# Lego-like Mixing and Matching of Engineered Bacteria Configure Full Subtractor and Adder Through an Artificial Neural Network Type Architecture

**DOI:** 10.1101/2023.06.15.545057

**Authors:** Deepro Bonnerjee, Saswata Chakraborty, Sangram Bagh

## Abstract

One of the long-term goals of synthetic bioengineering is to create configurable and programmable biological systems by just mixing and matching "LEGO"-like bio-modules. Here, we introduce a configurable and modular multi-cellular system where, from a small library of nine discrete engineered bacterial cells, a full subtractor and a full adder can be built on demand by just mixing and matching seven appropriate cell types in a culture. Here, each set of engineered bacteria was modelled as an ‘artificial neuro-synapse’ that, in a co-culture, formed a single layer artificial neural network (ANN) type architecture that worked as a biochemical full subtractor or full adder. The system is configurable with interchangeable cellular modules, whereby through simply interchanging two cell types in the subtractor culture, a full adder can be built and vice versa. This Lego-like mix and match system is mathematically predictive, and provide a flexible and scalable means to build complex cellular functions. This work may have significance in biocomputer technology development, multi-cellular synthetic biology, and cellular hardware for ANN.

## Introduction

Performing computations with engineered living cells has enormous importance in biocomputer technology development at the micron scale [1-3], where microprocessor-based computers have limitations due to energy, cost, and technological constraints [3]. However, the success of biocomputing technology depends on expanding the horizons of implementing complex computational device functions in living cells. Such system designs [4-6] follow the hierarchical electronic design principle, where the component genetic logic gates are layered according to the electronic analogous design either in a single cell [7-9] or in a multicellular distributed format [10-12]. However, layering multiple logic gates in a single cell result in crosstalk among various parts and hinders building complex functionality [11, 12]. Multicellular distributed computing has been adopted to build more complex functions, where various engineered cells carry a part of the integrated circuits and are connected with other cells through engineered chemical communications [10-12]. As each engineered cell is physically insulated from the other engineered cells, this reduces the crosstalk. Such strategies were used to build integrated logic circuits [10] and maze-solving biocomputers[13] in bacteria and an adder [11] and analog-to-digital converter[12] in mammalian cells.

However, such systems are complicated to build, require multiple layers of various engineered cells, and depend on the availability of the orthogonal diffusible chemical communication channels. Further, they are nonconfigurable, mathematically unpredictable, and demonstrate a single function. One of the long-term goals of synthetic biology and bioengineering is to create configurable and programmable biological systems by just mixing and matching "Lego"-like bio modules to build multiple predictable biological functions [14].

Here we introduce a configurable and modular engineered multi-cellular system where, from a small library of nine discrete engineered bacterial cells, a full subtractor and a full adder can be built on demand by just mixing and matching seven appropriate cell types in a culture. The system is configurable with interchangeable cellular modules, whereby by simply interchanging two cell types, a full subtractor culture can be reconfigured into that of a full adder and vice versa.

A full subtractor and a full adder form indispensable parts of modern ICs where they are found in the arithmetic logic units and the digital signal processing units to perform binary subtraction and binary addition respectively. A full subtractor is a function that performs subtraction of two bits, one is minuend and other is subtrahend, taking into account borrow of the previous adjacent lower minuend bit. The digit from which another digit is to be subtracted is known as minuend. Similarly, the number which is subtracted from the minuend is known as a subtrahend. It has three inputs, the minuend, subtrahend, and borrow-in from a previous full subtractor and two outputs, the difference and output borrow, respectively. The output borrow is connected as the input of a subsequent full subtractor. Full Adder is a function that adds three input bits and produces two outputs. The first two inputs get added along with a third input, that represents an input carry from a previous full adder. The outputs are the sum and a carry-out to the next adder. The output carry of a first adder is connected to the input of the second adder. Both the functions are complex to build with cells. No cellular subtractor has been realised yet. A mammalian multicellular full adder [11] required twenty-two component genetic logic circuits in nine different types of cells, which were layered according to the electronic circuit design and connected through engineered cellular communications.

To create two complex and challenging functions in an easy, configurable Lego-like format, we adapted the basic concepts of artificial neural network (ANN) architecture to represent engineered bacteria as artificial neuro-synapses or “bactoneurons” wherein a hetergenous population of such cells constitute a single layer ANN. Previously, we demonstrated [15,16] that such single-layer ANN-type architectures with engineered bacteria could generate multiple computing functions like decoders, encoders, majority functions, Fredkin gates, and double Feynman gates.

Motivated by the success of this design concept, in this work, we built two single-layered Bactoneural-Networks (BNNs), each comprised of 7 bactoneruons formulated from a library of a just 9 bactoneuron types, which can implement two complex computing functions, a full subtractor and a full adder. The two BNN’s differed from each other by 2 bactoneuron types and a simple re-configuration effected by interchanging these two, reprograms the entire bactoneural layer between two separate but equally complex computing functions. The reconfigurable full subtractor and adder made from engineered bacteria have several advantages: i) the system is configurable and the cells are interchangeable. It showed two independent functionalities by replacing just two bactoneurons: ii) the Lego-like wetware design is modular, simpler, and smaller, the system creates two complex and challenging functions in an easy format; iii) both the subtractor and adder were realised by 7 different cell populations arranged in a single layer, without the need of layering the cells by engineered chemical communications; iii) the behaviour of each module and the whole system can be described with mathematical equations of a single type, and each cellular type showed agreement between simulation and experiments; iv) robust rules for engineering such systems is in place; v) the design is general and can be built with any type of cells.

## Results

### Design of the configurable subtractor and adder in ANN type architecture

First, the subtractor and adder truth tables are mapped in a single-layer ANN-type architecture format (Fig. 1, Fig. S1) following a set of rules coherent with ANN[15]. In short, first we segregated each of the truth tables into its individual output channels and considered the relationship of that specific output with all the input combinations. Next, we grouped those combinations into smaller truth tables such that each input corresponding to a particular output possess a weight with only one unique type of sign: ‘+," "-," or "0’. We now reduced the resultant truth table by ignoring any component with a weight value of "0’. If the output value 1 (true) in the smaller resultant truth table appeared only once, we terminated fragmenting the truth table. Otherwise, we kept fragmenting the truth table until the above condition appeared. The nine resultant truth tables represented the functional and mathematical characteristics of nine "artificial neuro-synapses" (Figure 1a), or unit bactoneurons, which need to be engineered as cellular devices. Together, the nine bactoneurons work as a configurable cellular system, such that when the corresponding seven unit bactoneurons are assembled into a single layered ANN-type architecture, both the subtractor and adder could be realised by interchanging two bactoneurons from a library of nine.

**Figure 1:**
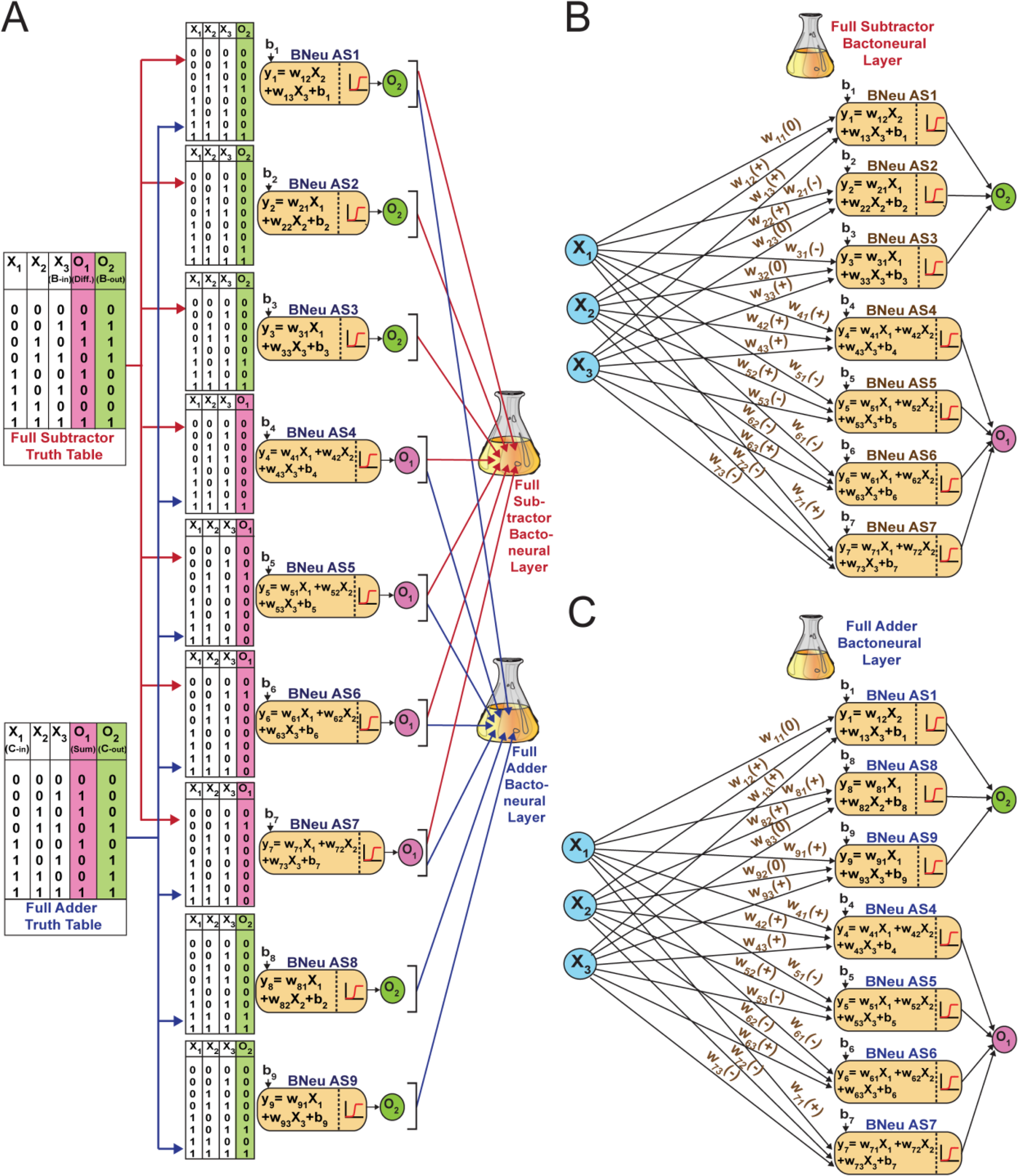
Fragmentation and mapping of the truth tables of Full Subtractor & Full Adder into configurable bactoneural layers. **(A)** The two functions Full Subtractor (above) and Full-adder (below) are first fragmented into smaller simpler functions according to the rules of ANN and are represented by the vertical array of sub-truth tables. The sub-functions generating out of Full Subtractor are indicated through red connecting arrows and the ones generating out Full Adder are shown through blue connecting arrows. The corresponding output channels are shown in colour (O_1_ is magenta and O_2_ is green). These functions get mapped onto the Bactoneurons where the parameters of the summation function are implemented through and emanate from synthetic genetic circuits fabricated based on established design principles connecting ANN and transcriptional regulation. X_1_, X_2_ & X_3_ represent the input channels, w_ij_ represents the weight of the j^th^ input for the i^th^ bactoneuron. b_i_ represents the Bias of the i^th^ bactoneuron. The weighted summation function activates the activation function of the bactoneuron that respond through expression of fluorescent proteins that serve as output channels. Each bactoneuron is connected to any one output channel. Finally, populations of appropriate bactoneurons can be mixed and matched in a heterogenous culture to constitute bactoneural layers that can be configured according to the function desired. **(B)** Configuration of the bactoneural single layer network for performing Full Subtractor function. The bactoneural layer composed 7 bactoneurons (AS-1-7) takes weighted inputs, combines them linearly in a summation function and responds through the output channels after activation through a sigmoidal function. **(C)** Configuration of the Full Adder bactoneural layer composed of Bactoneurons AS-1,4,5,6,7,8,9. The figure demonstrates how reconfiguration of the bactoneural layer by replacing just 2 bactoneurons out of seven, AS-2 & 3 with AS-8 & 9, reprograms the system from Full Subtractor to Full Adder.

### Design of cellular devices and unit bactoneuron

The bactoneurons were constructed by building transcriptional synthetic genetic circuits in the bacteria (Fig. 2). The inputs X^1^, X_2_, and X_3_ were replaced with three chemical inducers, N-acyl homoserine lactone (AHL) isopropyl b-D-1 thiogalactopyranoside (IPTG), and anhydrotetracycline (aTc) respectively. The outputs O_1_ and O_2_ were manifested with fluorescent proteins E2 crimson, and EGFP respectively (Figure 2). Synthetic genetic circuits inside bacteria process chemical inputs and produce output by expressing fluorescent proteins, such that the input-output relation mimics a conventional activation function[17] for an artificial neuro-synapse (eq 1). Thus, for a given bactoneuron j,

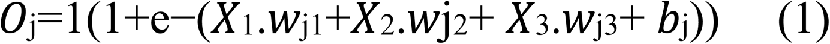

**Figure 2:**
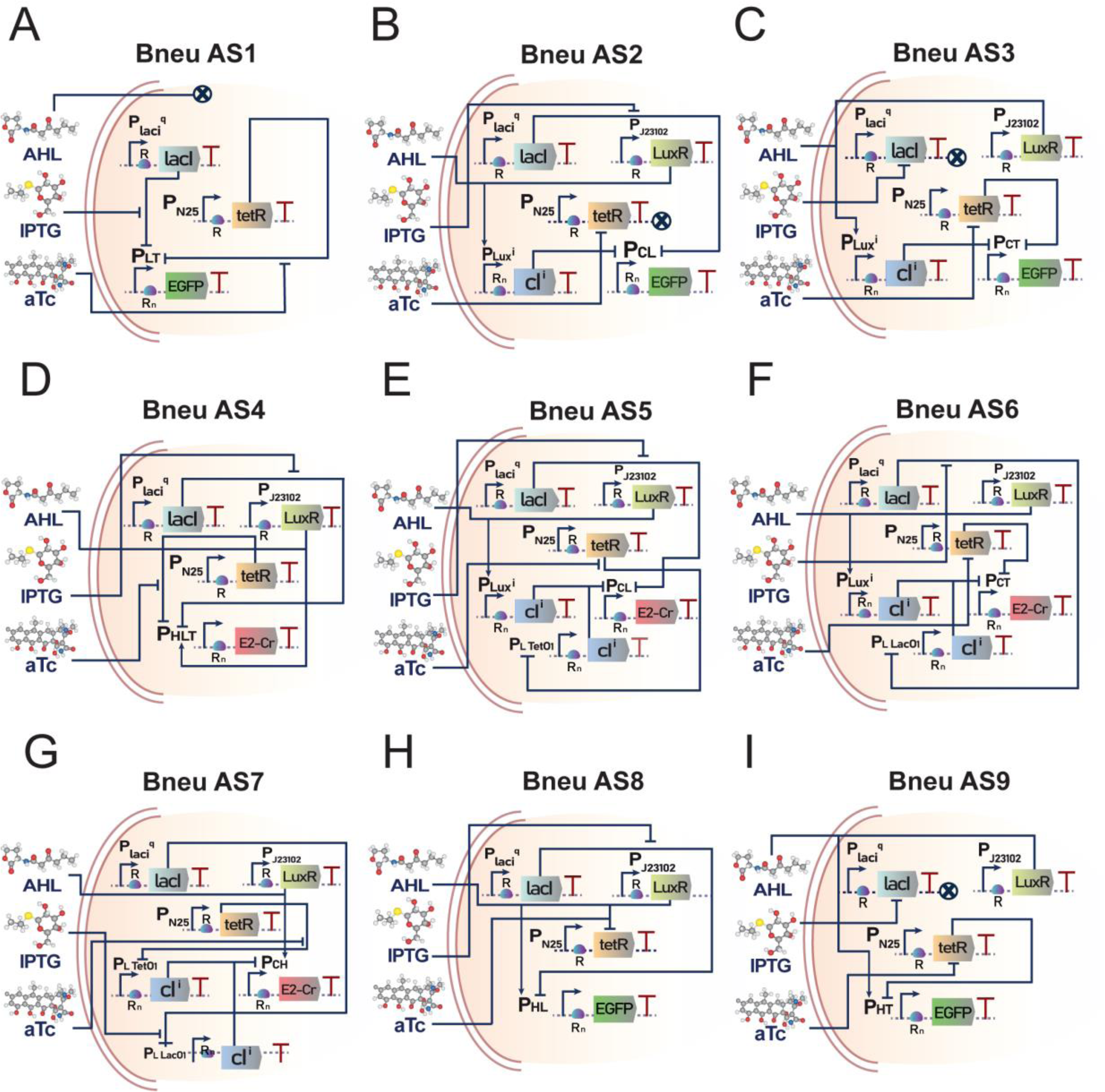
Generic maps of the synthetic genetic networks that encode corresponding specific functions to each bactoneuron type. **(A)** Bneu AS1, **(B)** Bneu AS2, **(C)** Bneu AS3, **(D)** Bneu AS4, **(E)** Bneu AS5, **(F)** Bneu AS6, **(G)** Bneu AS7, **(H)** Bneu AS8 and **(I)** Bneu AS9. The three abstract inputs X_1_, X_2_ & X_3_ are served by extracellularly provided AHL, IPTG & aTc respectively. Abstract outputs O_1_ and O_2_ are served by E2Crimson & EGFP respectively. In different combinations of the three inputs, they influence the transcriptional machinery of the synthetic genetic circuitry by interacting with transcription factors LuxR (activator), LacI & TetR (repressors) respectively by either activating the factor (LuxR) or deactivating it (LacI & tetR) through conformational changes. This interaction affects the transcription factor’s binding affinity with its respective operator sites on the sequence of various promoters within the genetic circuit in a non-linear fashion (activation by LuxR or repression by LacO1 or TetO2) thereby enabling the implementation of the activation functions. R refers to RBSs that were remained constant and was not part of iterative updates. R_n_ refers to RBSs that were part for the iterative process where “n” corresponds to RBS number. cI^i^ refers to either of 2 variants of the lambda repressor proteins cI^w^ or cI* which were interchanged as part of iterations. cI^w^ refers to the wild type version of cI whereas cI* refers to the frameshifted version^18^. Plux^i^ refers to promoters Plux^w^ or Plux* which were interchanged across iterations. Gene expression cassettes that do not have any downstream effect on the output and are associated with “0” weight inputs are shown with their interaction lines terminating in a **“X”** sign.

where O_j_ is the fluorescent protein output of the bactoneuron, X_i_ represents the input inducer concentration, w_ji_ represents the weight of the corresponding input X_i_, and b_j_ represents the bias for the bactoneuron j. We designed the molecular-genetic circuits of all the bactoneurons based on the signs of the ‘weight’ parameter in activation function equations for the corresponding bactoneuron, (Fig. 2). The positive and negative weight values in an activation function of a bactoneuron respectively correspond to the activation and repression of output reporter gene expression mediated by a specific inducer. Zero weight corresponds to an invariant relation.

The transcriptional genetic circuits (Fig 2) were built based on a set of synthetic promoters on which transcription factors LuxR, LacR, TetR, and Ci [18] bind to either individually or simultaneously to hinder and/or activate the transcriptions. The transcription factors, promoters, and their interactions formed various feed-forward, feedback, and combination mechanisms based on the bactoneuron circuit designs (Fig. 2). The mode of transcription was controlled by the interactions of the transcription factors with appropriate input chemicals (AHL, IPTG, and aTc) through allosteric regulation. The quantitative relation between inputs and outputs within a proper parameter range should demonstrate appropriate bactoneuron behaviour.

### Construction, characterization, and optimization of bactoneurons

Among the nine bactoneurons required to configure a subtractor or adder, the basic form for five two-input bactoneurons (BNeu AS1, AS2, AS3, AS8, and AS9) were available from a bactoneuron library we made earlier [15,16]. The rest of the four bactoneurons (BNeu AS4, AS5, AS6, and AS7) are three input systems that were built from scratch. We first constructed the initial set of molecular constructs according to the design (Fig. 2) and inserted them into DH5αZ1 strains [19] of *E.coli*. Next, we measured fluorescence protein expression at ‘0’ and ‘1’ (saturated) inducer concentrations (Figs. S2, S3, S4, and S5). These experiments gave us fold changes between the highest output signal (value "1") (F.C.) and the highest leakage, (where the leakage was defined as the basal level expression of the reporter fluorescent proteins under the input conditions, where output expression should be zero) total leakage from all possible inducer (0,1) combinations (Σ leakage, Σ L.), and variation in the highest signals among various bactoneurons (S.V). Further, we performed dose response experiments as a function of various inducer concentrations (Figs S2-S5) by varying one inducer concentration while keeping the others as constants (saturated or ‘0’ concentration, as the case may be). The dose response behaviours were fitted with the activation function equation (equation 1), and the weight and bias parameters were estimated from the fitted equations (Figs. S2-S5). The initial constructs for AS5, AS6, and AS7 showed suboptimal behaviour. They (AS5A, AS6A, and AS7A) generally had i) low fold changes (F.C.) between the highest output signal (value "1") and the highest leakage, ii) high total leakage (Σ L.), iii) low weight values, and iv) high variability of the signals at ON states among various bactoneurons (S.V.) (Fig 3C). We defined/fixed a set of optimum values arbitrarily or based on previous observations [13,15] (Fig. 3A). The minimum fold change between the highest leakage and the ON state was set at 7. The maximum total leakage was set at 0.2 and 0.4 for 2-input and 3-input bactoneurons, respectively. The weight values define the sharpness of the transition from the OFF state to the ON state as a function of input. The minimum weight values were set as 8.5, 9.5, and 10.5 for input AHL, IPTG, and aTc, respectively. Further, we measured the signal strength at ON states of all bactoneurons expressing a single fluorescent protein (EGFP or E2-Crimson) and fixed the maximum variation to be within 20%. Though bactoneuron AS4 satisfied the optimal set points in its first construct, the initial constructs for AS5, AS6, and AS7 failed. The measurements performed on all the two input bactoneurons taken from the library (BNeu AS1, AS2, AS3, AS8, and AS9) showed optimum behaviour.

**Figure3:**
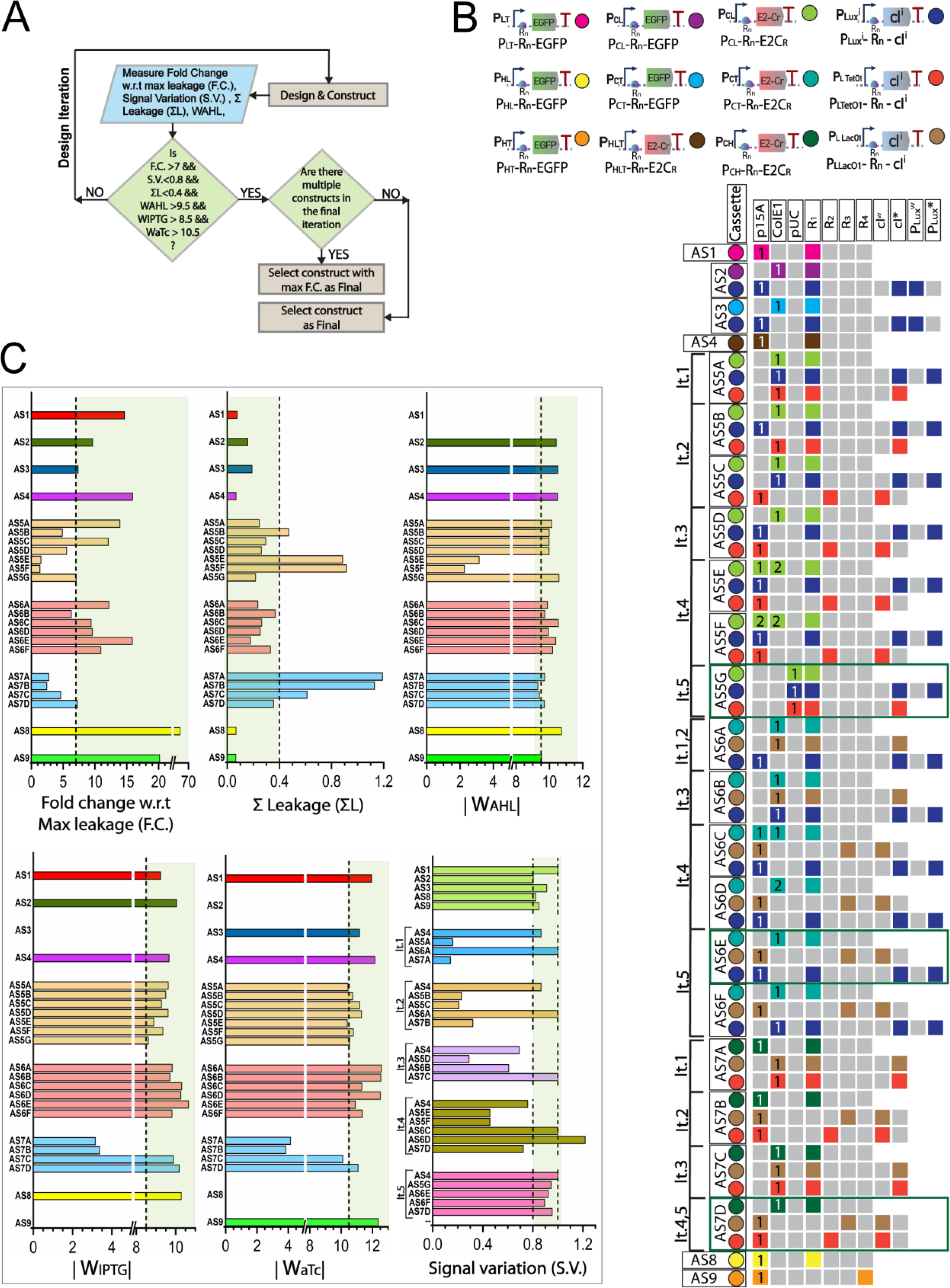
Details of the process of iterations to optimize Bactoneuron parameters. **(A)** Flowchart of the algorithm followed for bactoneural parameter optimization iterations. **(B)** The bioparts configuration space and details of the iterations to update bactoneural parameters. Biological designs of each bactoneuron construct involves one or more candidate gene expression cassettes that were chosen for design configuration iterations. Such cassettes are grouped with colour-coded circles under the construct name. Biological designs were iterated through updating either the relative copy number of a cassette (high copy pUC or medium copy ColE1 or low copy p15A) and/or by modulation of the translation initiation rate by changing the cassette’s RBS strength (R_1_, R_2_, R_3_ or R_4_) and/or by modulation of the strength of repression by lambda repressor by altering between wild type cI^w^ or mutated cI* and/or by modulating the promoter strength of the AHL+LuxR activated promoter Plux^i^ by altering between either of two variants Plux^w^ & Plux*. The specific bio-part configuration used for the gene expression cassette in question for an iteration is indicated through squares that glow with the same colour as of the cassette colour code. Grey square indicates an unused configuration. For bactoneurons that went through multiple design iterations, the corresponding constructs generated at each iteration are grouped under its iteration number denoted by “It.n” where n indicates iteration number. Constructs that were chosen as final are highlighted in green boxes. Bactoneuron contructs under no iteration bracket denotes no design iteration performed for them. **(C)** Parameter values achieved by different bactoneuron constructs. The set threshold and acceptable regions for each parameter are indicated by the dashed lines and the light green shaded area respectively. “Signal variation” (S.V.) is a comparative parameter drawn by scaling the fluorescence signal (@ “ON” state) of all bactoneurons in an iteration set (with the same output channel) by considering the strongest signal of all bactoneurons as “1”. Thus, S.V. is shown in iteration groups.

We performed a detailed failure analysis on the disqualified bactoneuron constructs (Table S1). Now, keeping the general molecular design framework intact (Fig. 2), we optimised those suboptimal bactoneurons by modifying the genetic constructs, altering the interaction strength among the various transcription factors and promoters, and modulating their relative amounts in the cell. This had been done by changing plasmid copy numbers, using mutated forms of transcription factors, altering ribosome binding sites and promoter sequences, and redistributing the bioparts (Fig. 3B). Guided by the optimisation flow chart (Fig. 3A) and the failure analysis (Table S1), the optimisation was conducted in multiple iterations (Fig. 3B) until quantitative set points were reached (Fig. 3C). The schematics of genetic constructs and bioparts distribution among various plasmids in each iteration for each bactoneurons are shown in figures S6–S9.

### Simulation and validation of optimized bactoneurons

Based on the fitted parameters for each optimised bactoneurons, we performed computational simulations using parameterized activation functions. (Fig. 4). Further, we experimentally tested the behaviour of each optimised bactoneurons by simultaneously changing the concentrations of combinations of two inputs from among AHL, IPTG, and aTc and measuring the fluorescence at each data point. To reduce the experimental bias, in each case, a different set of inducer concentrations was selected compared to the dose response experiments (Table S2). These validation tests showed a close topological match between the simulated and experimental behaviour (Fig. 4).

**Figure 4:**
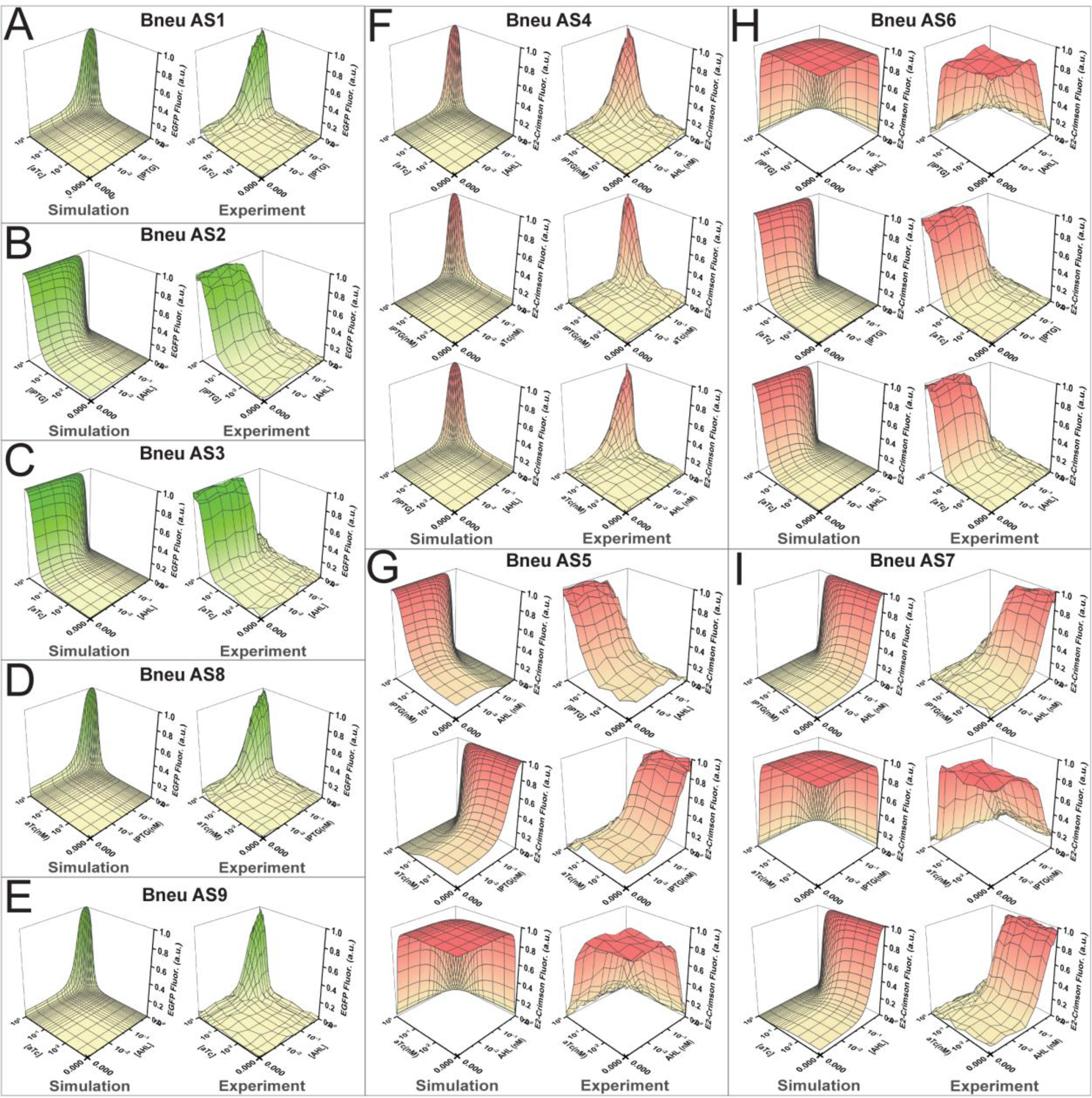
Simulations drawn from parameterized mathematical models of optimized Bactoneurons and its validation through validation experiments. Mathematical models were developed from parameters obtained from Dose response experiments (see Supplementary figures S2-5) and simulations were developed from these models by varying 2 inducers across 50x50 concentration points (while keeping the third inducer, if any, constant at “0” or “1” as the case may be) (see Methods) and their response in corresponding mathematical output signal intensity for each point. The simulations are shown towards the left of a panel for a bactoneuron. These simulations were validated by finding a good topological match with the validation experiments where the same corresponding set of inducers were varied across 16x16 concentration points in an actual experiment (that were different from those used for dose-response, see Supplementary table ST2) keeping the third, if any, constant, and plot with the corresponding reporter protein fluorescence. The validation experiments are shown towards the right of a panel for a bactoneuron. Panel **(A)** for Bneu AS1, **(B)** Bneu AS2, **(C)** Bneu AS3, **(D)** Bneu AS8, **(E)** Bneu AS9, **(F)** Bneu AS4, **(G)** Bneu AS5, **(H)** Bneu AS6 and **(I)** Bneu AS7. Bactoneurons connected to an EGFP output channel are represented in green shaded topology and those connected E2Crimson are shown in red shaded topology.

**Figure 5:**
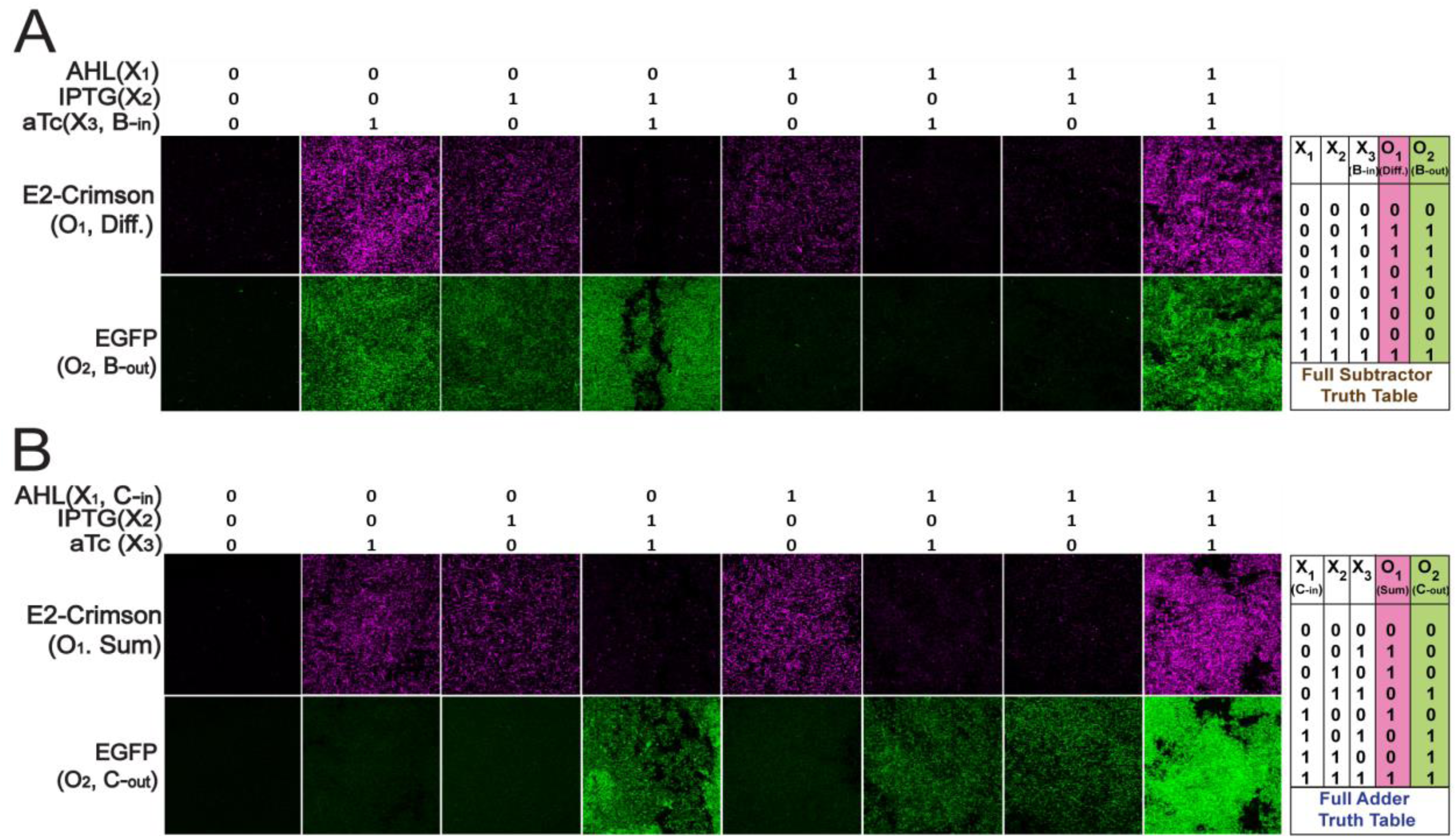
Fluorescence microscopy for A) Full Subtractor and B) Full Adder. Combinations of appropriate Bactoneuron types as required to constitute the corresponding Bactoneural layer were co-cultured for 16hours (with a sub-culture of 100 times dilution at the 10hour time point) with appropriate inducer combinations as per the respective truth table (shown to the right). The cultures were PBS washed and viewed as slides under relevant laser channels and emission filters (see methods). Heterogenous populations of E-coli cells forming the bactoneural layer can be seen in the fields under DIC/merged channels (see supplementary figure S10) where they respond in accordance to the input combinations. Each sub-population of bactoneurons in the bactoneural layer either gets activated (indicated by fluorescence from its respective output protein) or remains inactive (showing no fluorescence) based on the input inducer combinations. Logical adherence to the respective truth tables can be observed indicating proper operation of the Full Subtractor or Full Adder functions through the bactoneural layer.

### Mix and match bactoneurons for a configurable adder and subtractor

The nine optimised bactoneurons now worked as a configurable cellular system that could generate a full subtractor or a full adder. By mixing BNeu AS1, AS2, AS3, AS4, AS5, AS6, and AS7 in a co-culture, we created a bactoneural layer that harboured the full subtractor function. Next, the subtractor co-culture was exposed to various combinations of input chemicals (AHL, IPTG, and aTc) according to the truth table, and the output fluorescence was seen with a fluorescence microscope. For the Full Subtractor, aTc served as the input Borrow (Bin), AHL served as the minuend and IPTG serves as the subtrahend. The E2-crimson expression (Output O_1_) signified difference, and the EGFP expression (O_2_) signified borrow-out. The results showed proper subtractor behaviour (Fig. 4A; Fig. S10A). Similarly, by just replacing BNeu AS2 and AS3 with BNeu AS8 and AS9 we could re-configure the bactoneural layer for a full adder behaviour, where aTc and IPTG served as the general inputs and AHL served as the input Carry (Cin); E2-Crimson expression signified sum and EGFP expression signified carry-out. The experimental results showed proper adder behaviour under fluorescence microscope. (Fig. 4B, Fig. S10B).

## Discussion

We demonstrated a configurable full subtractor and a full adder using engineered bacteria. The system is modular, whereby mixing and matching seven different bacterial populations can give rise to either a subtractor or an adder. Out of seven, five of the modular bacteria (BNeu AS1, AS4, AS5, AS6, and AS7) are common in both subtractor and adder, and by replacing only two bacterial types, it can switch between either a full subtractor or full adder. The universal approximation theorem suggests that a feed-forward artificial neural network (ANN)-type architecture may approximate a wide variety of non-linear functions [17]. Thus, an ANN architecture might be a good strategy to build complex biocomputing functions. In this work, full subtractor and adder were created through a single-layer ANN-type architecture, where no layering of the cells with chemical communication was required. Thus, this design makes the construction of complex computing functions in cells easier. The absence of a second layer of cells with chemical communication might make the system respond faster to input signals.

Modularity and configurability are important hallmarks of engineering systems [20]. This helps to manage the complexity, perform parallel work, and accommodate future uncertainty. In this work, each engineered cell, which represented an artificial neuro-synapse within an ANN framework, worked as a modular and configurable system such that mixing and matching could generate two different functions. A few of the bactoneurons were taken from previous works, where they worked as modules to generate other functions. Interestingly, those ‘bactoneurons’ worked perfectly well within the configurable subtractor and adder system, suggesting the subtractor and adder possess a high degree of modularity and scalability.

Complex decision-making capability at the micron scale may have unprecedented applications in smart therapy [21-23], bioproduction [24] and bio-remediation [25]. Theoretical studies [26-28] predicted that to perform complex and large computations, multicellular distributed computing was necessary for its scalability, error tolerance and burden reduction per individual cell. However, often such experimental systems are fragile and unpredictive [3]. One of the hurdles in building functional multi-cellular systems is the unavailability of quantitative parameters, which are required for understanding as well as controlling the system [29-31]. In this work, each of the modular bactoneurons was quantitatively characterised and mathematically predictable. As the whole ANN architecture related to its modules (nodes) through a mathematical framework, the behaviour of the ultimate function, subtractor or adder, was also predictable.

Though the work has been demonstrated with bacteria, the design of the ANN architecture and its individual ‘bactoneurons’ is abstract and general. Thus, it can be implemented with any type of cell, bacterial or mammalian.

Taken together, this ‘lego-like’’ mix and match modular and configurable cellular system provide flexible and scalable means to build complex biological functions with engineered cells. This work may have significance in biocomputer technology development, multi-cellular synthetic biology, and cellular hardware for ANN with potential industrial and therapeutic applications.

## Materials and Methods

### RBS design and promoters, genes, and plasmids

Promoters, ribosome biding sites (RBS), genes & transcription terminators were all assembled using the Network Brick pipeline [32] into pZ expression systems [19] according to the corresponding genetic circuit designs. RBS with various strengths were designed and translation initiation rates for designed RBSs (Table S3) were calculated thermodynamically from RBS calculator [33]. Standard molecular biology protocols were used. Restriction enzymes, T4 DNA ligase, buffers & DNA ladders were from New England BioLabs. PCR reactions were performed using the KOD Hotstart DNA polymerase (Merck Millipore). Plasmid isolation, PCR purification & Gel extraction were done using QIAGEN kits. All promoter sequences, gene sequences and primer sequences are shown in table S4. Details of the cellular devices and plasmids are shown in figures S6-S9. Primers were synthesized by Integrated DNA Technologies, Singapore. Oligos & gene Products were synthesized by Invitrogen GeneArt Gene Synthesis service, Thermo Fischer. All constructs & bio-parts were sequence verified by Eurofins Genomics India Pvt. Ltd., Bangalore, India.

### Bacterial strains and culture conditions

For all cloning purposes *Escherichia coli* DH5α strain was used. For all experiments, DH5αZ1 strain [18] was used as the chassis organism. Chemically competent DH5αZ1 were co-transformed with sequence verified plasmid constructs as per design of ach of the bactoneurons and were plated onto LB-agar, Miller (Himedia) plates with appropriate antibiotics (ampicillin (Himedia) at 100 µg/ml, chloramphenicol (Himedia) at 34 µg/ml). For all liquid cultures, LB-broth, Miller (Himedia) was used with appropriate antibiotics. Appropriate chemical inducers were provided into liquid cultures (AHL (C_10_H_15_NO_4,_) (Merck) 0 – 5000 nM, IPTG (Himedia) 0- 10,000 µM, aTc (Sigma) 0 -200 ng/ml. Incubation was done 37° C, ∼ 280 rpm in New Brunswick Scientific, Innova 42 incubator shaker.

### Testing of Bactoneuron constructs for adherence to respective truth-tables

Four distinct single colonies were picked from the plates and grown in liquid culture. For induction, overnight cultures were diluted (100 times) in fresh LB with appropriate inducer combinations as per the truth table (Fig. 1). This induction was given for a total of 16 h with a sub-culture of 1/100 dilution at the 10hr time-point. The ‘0’ and ‘1’ states signified zero and saturated inducer concentration.

### Dose Response Experiments

3 or 4 single colony-overnight cultures for each bactoneurons were set up in in LB-Broth as before. For an experimental set, each of these cultures were subjected to 16hour induction (with a sub-culture of 1/100 dilution into identical composition media at the 10hr time-point) in separate 3mL LB broth cultures with different inducer concentration points, where one inducer was varied across 10 or more intermediate concentration points (Table S2) between its “0” to “1” state (see scaling below) while the other inducer(s) was/were kept constant at “0” or “1” concentration as the case maybe. This was performed for each bactoneuron for each of the 3 or 2 inducers with non-negative weights.

### Fluorescence Measurements, Normalization and Scaling

Post 16hr induction cells were washed and re-suspended in phosphate buffered saline (PBS, pH 7.4) for fluorescence measurements. Cells were re-suspended into PBS such that OD_600_ values were between 0.7-0.9. These cultures were then loaded onto a black flat-bottom 96-well plate (Greiner) & using a Synergy HTX Multi-Mode reader (Biotek Instruments, USA), OD_600_ & fluorescence was measured using appropriate combinations of excitation and emission filters. Cells were excited by a white light source that had been passed through an excitation filter of 485/20 nm band-pass for EGFP and 610/10nm band-pass for E2Crimson while emission was collected by 516/20 nm band-pass filter for EGFP at gain 60 & 610/10nm band-pass filter at gain 70 for E2Crimson. To collect the fluorescence and OD data at least 3 biological replicates had been considered for each condition. To normalize the raw fluorescence reading obtained, to the number cells in the well, the raw fluorescence values were divided by the respective OD_600_ values of each well. Auto-fluorescence was measured as average normalized fluorescence of the untransformed DH5αZ1 set (no plasmid set) and subtracted from the normalized fluorescence value of the experimental set. The resultant value was averaged across biological replicates to obtain the fluorescence intensity at an induction condition. The above process can be mathematically represented as follows:

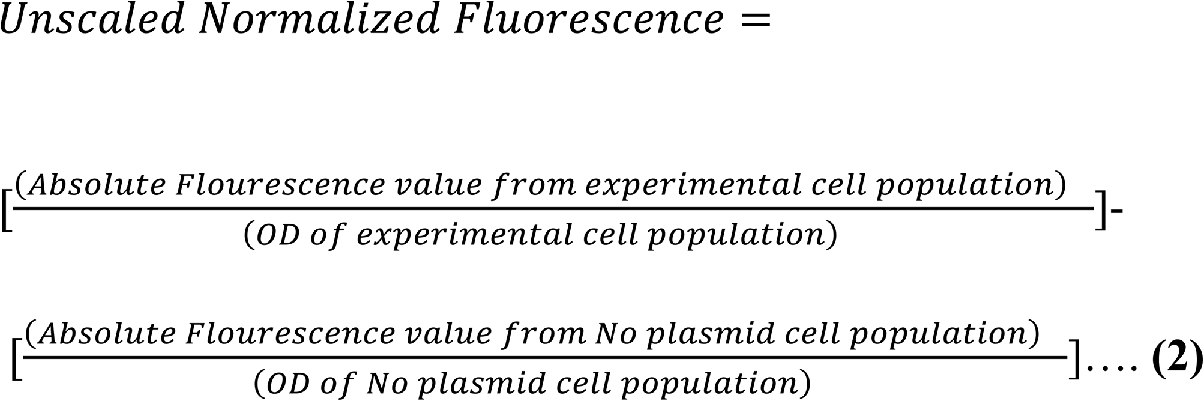

These normalized fluorescence values were then scaled down between 0 and 1 by considering the normalized fluorescence value at the induction point of maximum expected/observed fluorescence to be 1 and by dividing all other fluorescence values by this fluorescence value.

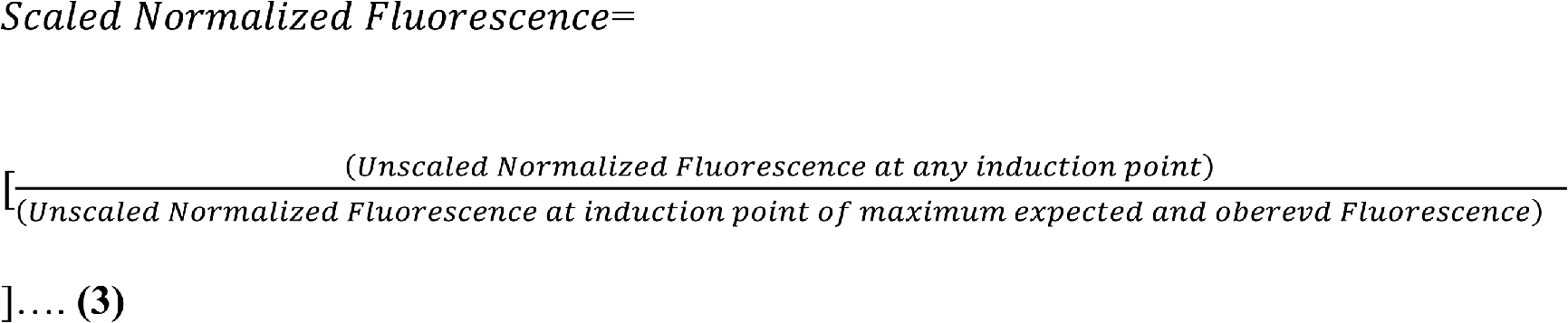

### Fitting & Scaling input concentration points

The scaled fluorescence values (y-axis) obtained were plotted against varying inducer concentration points (x-axis) onto a scatter plot on Origin2018 (OriginLab Corporation, USA). Using the in-built Orthogonal Distance Regression algorithm of the software, the scatter plot was fitted with a modified form of equation 1

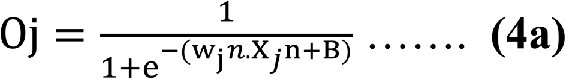

where, Y represents the normalized & scaled fluorescent outputs, W_jn_ represents the weight of the n^th^ inducer input for the j^th^ bactoneuron, X_jn_ represents scaled concentration for the n^th^ input and

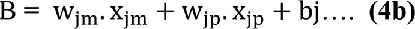

Where wj(m,p), xj(m,p) are weight and input for other two (m and p) inducers. B is a constant for an inducer specific dose response experiment for a specific bactoneuron. Inducer concentration points were scaled between 0 & 1 where “0” represents absence of the inducer in the experimental set & “1” represents that concentration of the inducer where Oj just attains a value of 0.999 for inducers with positive weight sign or 0.001 for those with negative weight sign. All values of concentration above this point were designated as “1”. The bias bj value of the bactoneuron was derived by solving from different parameterized versions of equation 4b from different dose response experiments.

### Simulation

All bactoneurons were modelled based on the activation function (equation 1). Parameterized activation function with numerical values of wj and bj were used for computational simulations for ach bactoneuron by generating 50x50 matrices of estimated output fluorescence values “*O*j” where 2 inputs simultaneously varied across 50x50 scaled relative concentration points between “0” & “1”. The third inducer (if any) was kept at “0” or “1” as the case may be. We used a fixed set of scaled relative concentration points to draw all the simulations as listed in table S5.

### Validatory experiments for verifying mathematical models & simulations

Verification of the mathematical models & simulations were done through validatory experiments subjecting an overnight culture of a single colony (n=1) of an appropriately transformed DH5αZ1 cell to 16hr induction as described previously, where 2 inducers were varied simultaneously across 16x16 concentration points (keeping the third inducer, if any, constant at “0” or “1” as the case may be) that were different from those that were used in the dose response experiments. The list of concentration points for validation experiments for different inducers & bactoneurons were are listed in table S2. Fluorescence readings were recorded, normalized, scaled and plot against the scaled concentration points of 2 inducers.

### Confocal Microscopy

Bactoneurons were grown overnight. Based on the bactoneural configuration of full subtractor or adder ANN (Figure 1), inoculum of each bactoneuron cultures were equivalently mixed and the combined inoculum was added with 100 times dilution into fresh LB broths with appropriate inducer combinations as per truth table. These 10hr co-cultures were again sub-cultured with 100 times dilution for another 6hours and the final co-cultures were washed twice with PBS (pH 7.4). The final washed cell pellets were again re-suspended in fresh PBS, added onto glass slides, covered with coverslips and sealed with a sealer. These were observed under 60X water immersion objective in a Nikon AIR si confocal microscope supplied with the resonant scanner and coherent CUBE diode laser system. The sample fields were subjected to excitation of the fluorescent proteins by sequentially illuminating with 2 laser channels, with λ = 488 nm for EGFP, and λ= 561 nm for E2-Crimson. Fluorescence emissions were captured through a T-PMT with relevant emission filters of 525/50 nm band-pass for EGFP & 700/75 nm band-pass for E2-Crimson. Differential interference contrast (DIC) images were also acquired for each captured field. ImageJ Fiji software was used for processing of acquired images.

## Author Contributions

D.B. and S.C. performed all the experiments. D.B. and S.B. designed the experiments, analyzed the data, and wrote the manuscript. S.B. conceived the study.

## Notes

The authors declare no competing financial interest.

## ACKNOWLEDGMENTS

This work was financially supported by Grants RSI4002 (Department of Atomic Energy, Govt. of India)

## Supplementary Information

**Figure S1:**
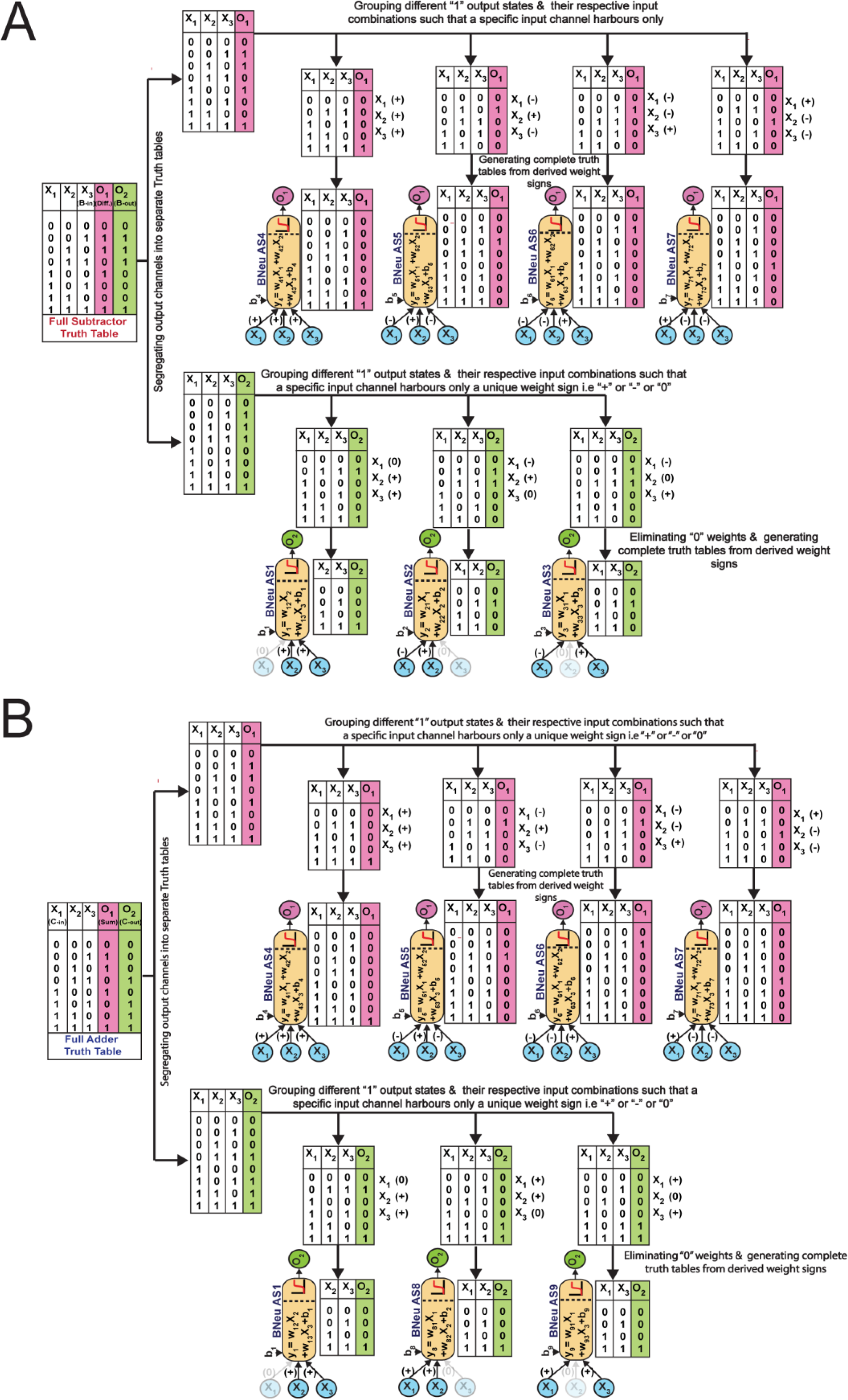
Fragmentation of Truth tables through ANN design rules and derivation of Bactoneurons with corresponding weight signs for A) Full Subtractor B) Full Adder. “0” weight inputs, if any, of a bactoneuron have been faded in the Bactoneural schematic.

**Figure S2:**
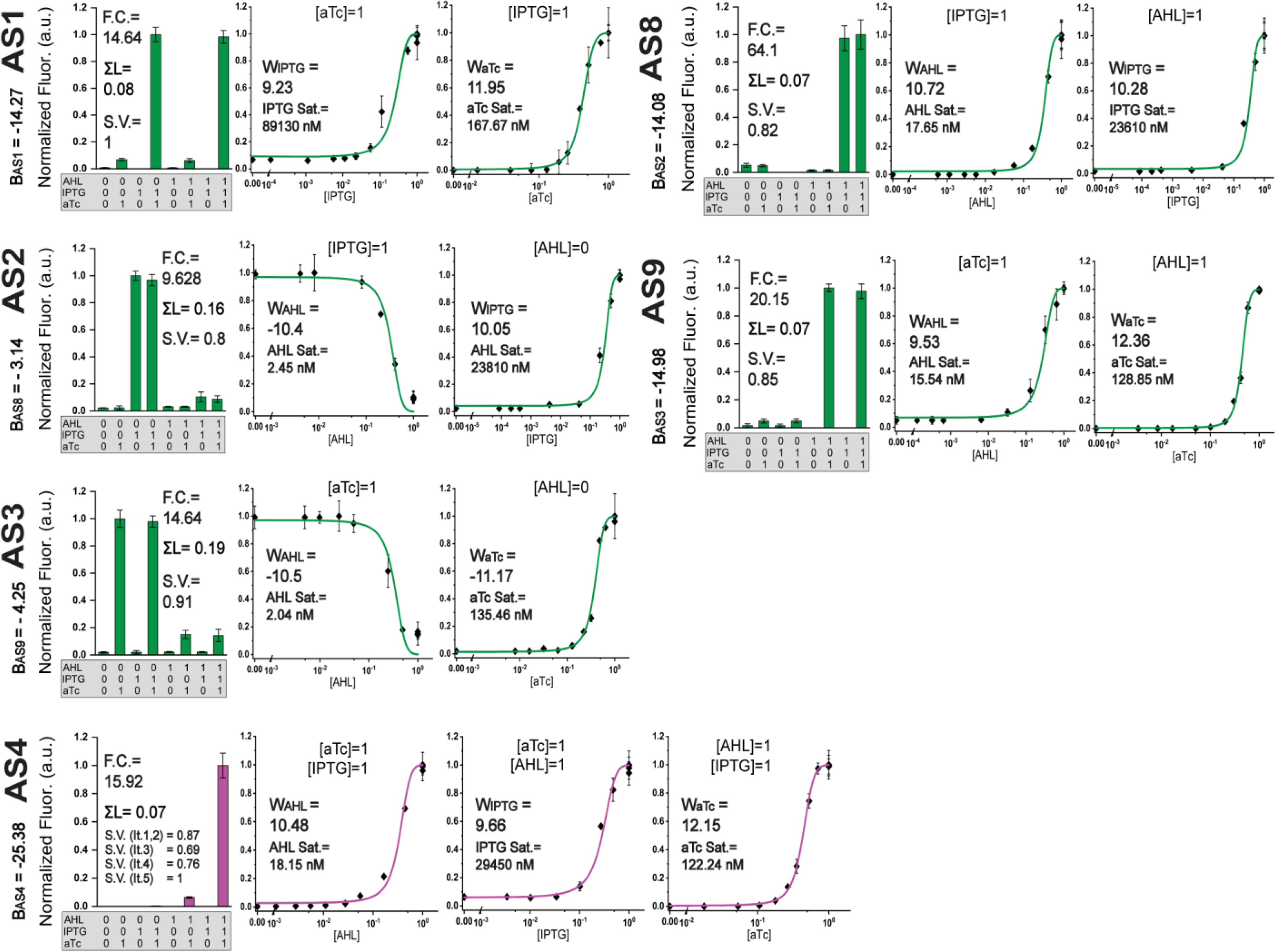
Details of characterization and Dose Responses experiments of Bactoneurons that went through no iteration and were selected as final. (AS1, AS2, AS3, AS4, AS8 & AS9). The bactoneurons were separately subjected to all possible combinations of input conditions (saturated concentration “1” or absent “0”) yielding corresponding parameter values of Fold Change (F.C.), Σ Leakage (ΣL) & Signal variation (S.V.) (shown in figure). “Signal variation” (S.V.) is a comparative parameter drawn by scaling the fluorescence signal (@ “ON” state) of all bactoneurons in an iteration set (with the same output channel) by considering the strongest signal of all bactoneurons as “1”. For Dose response, the concentration of one parameter was varied across various concentration points while the other two inducers were kept constant at “0” or “1” as appropriate (specified atop each dose response curve). The fluorescence readings obtained from the dose response experiments were fitted with the Log-Sigmoid function (see Methods) to derive the parameters Bias (B_AS#_) & Weights (W_inducer_) & the respective Inducer Saturation points (Sat.) were identified.

**Figure S3:**
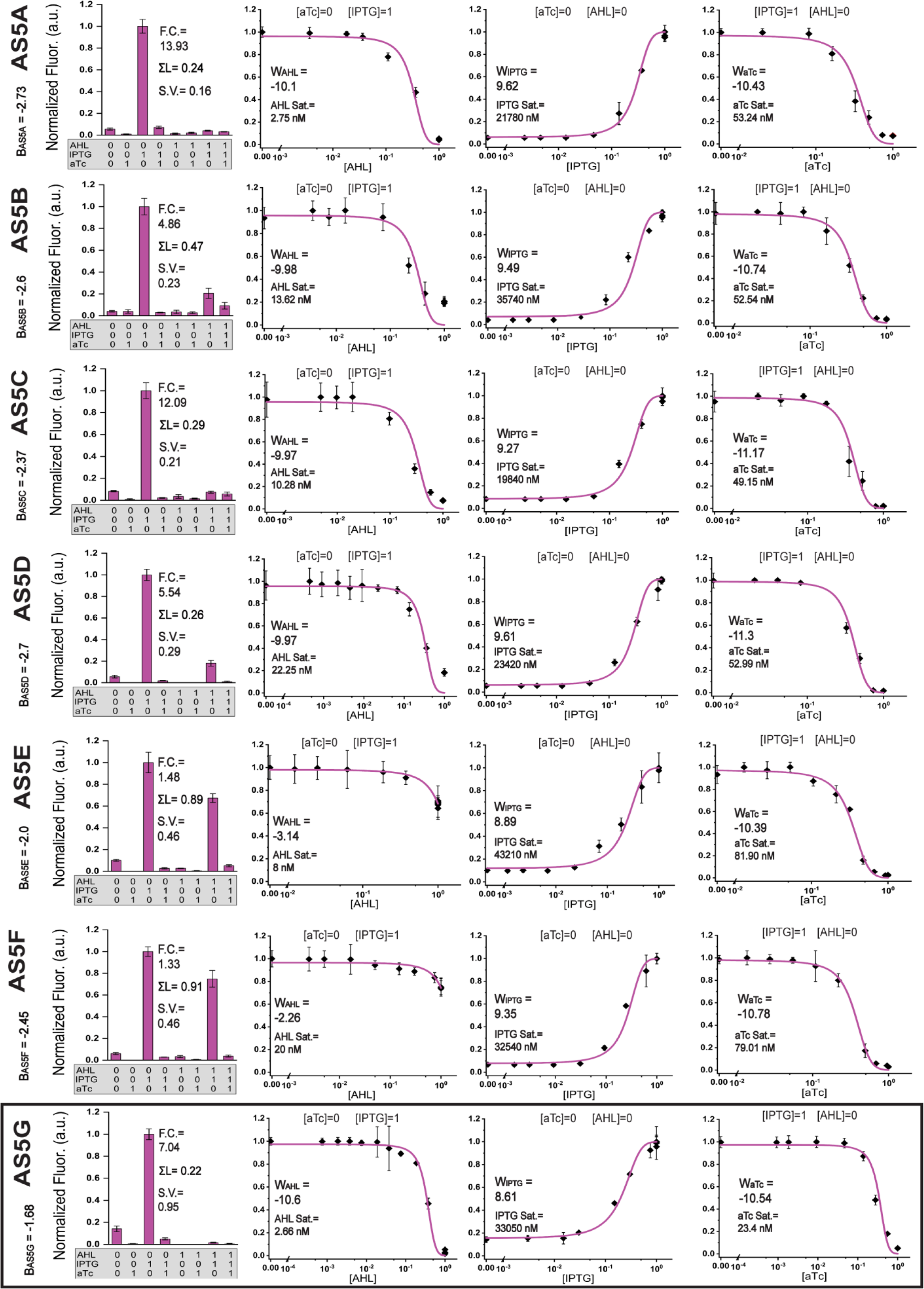
Details of characterization and Dose Responses experiments across iterations of Bactoneuron 5. (AS5A, AS5B, AS5C, AS5D, AS5E, AS5F & AS5G) yielding corresponding parameter values (shown in figure) of Fold Change (F.C.), Σ Leakage (ΣL), Signal variation (S.V.), Bias (B_AS#_), Weights (W_inducer_) & Inducer Saturation points (Sat.). The fluorescence readings obtained from the dose response experiments were fitted with the Log-Sigmoid function (see methods) to derive the Weight & Bias parameters. The induction state of the two constant inducers are specified atop the fitted curves. The final selected Construct is shown boxed.

**Figure S4:**
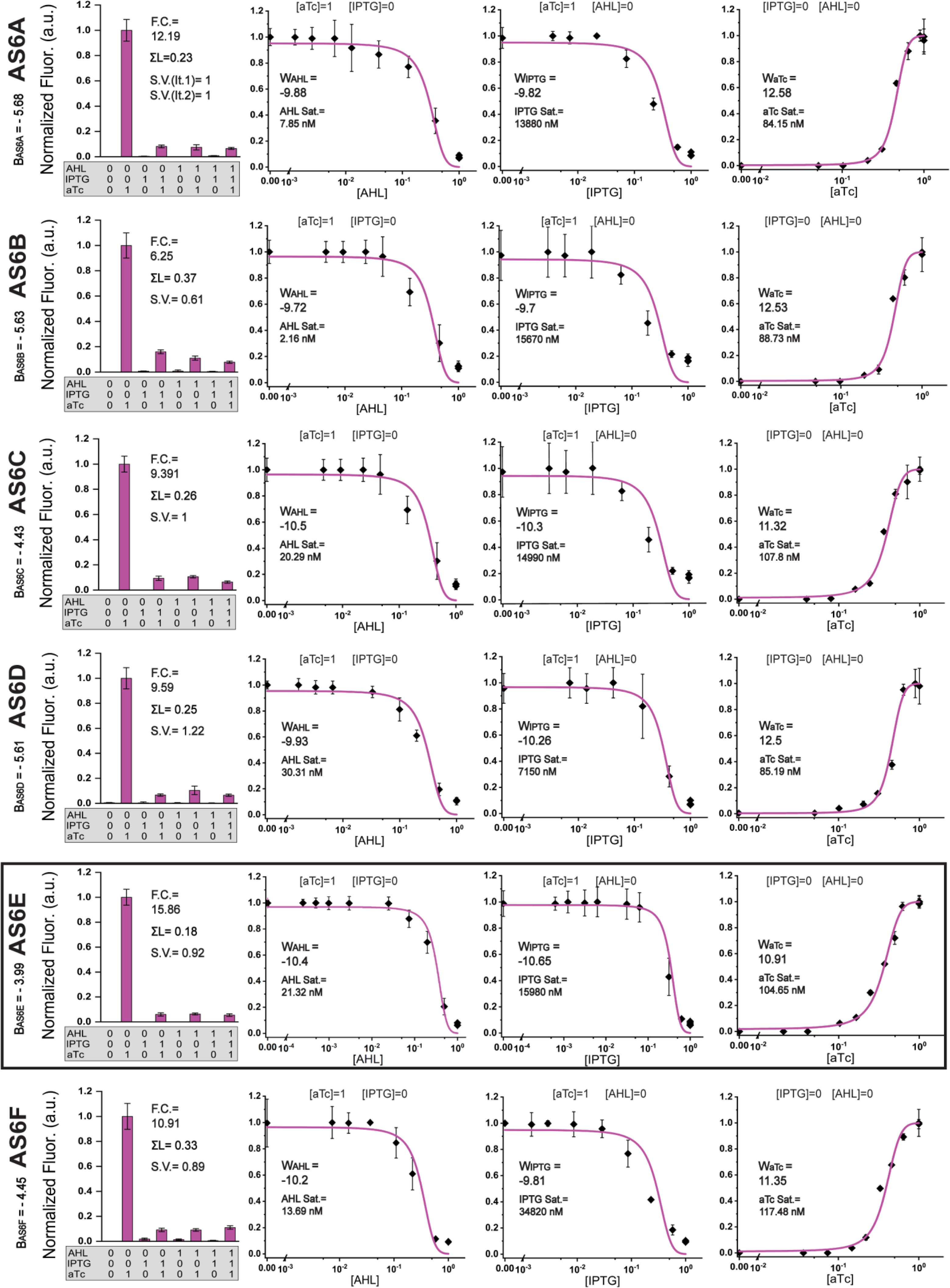
Details of characterization and Dose Responses experiments across iterations of Bactoneuron 6. (AS6A, AS6B, AS6C, AS6D, AS6E &AS6F) yielding corresponding parameter values (shown in figure) of Fold Change (F.C.), Σ Leakage (ΣL), Signal variation (S.V.), Bias (B_AS#_), Weights (W_Inducer)_ & Inducer Saturation points (Sat.). The fluorescence readings obtained from the dose response experiments were fitted with the Log-Sigmoid function (see Methods) to derive the Weight & Bias parameters. The induction state of the two constant inducers are specified atop the fitted curves. The final selected Construct is shown boxed.

**Figure S5:**
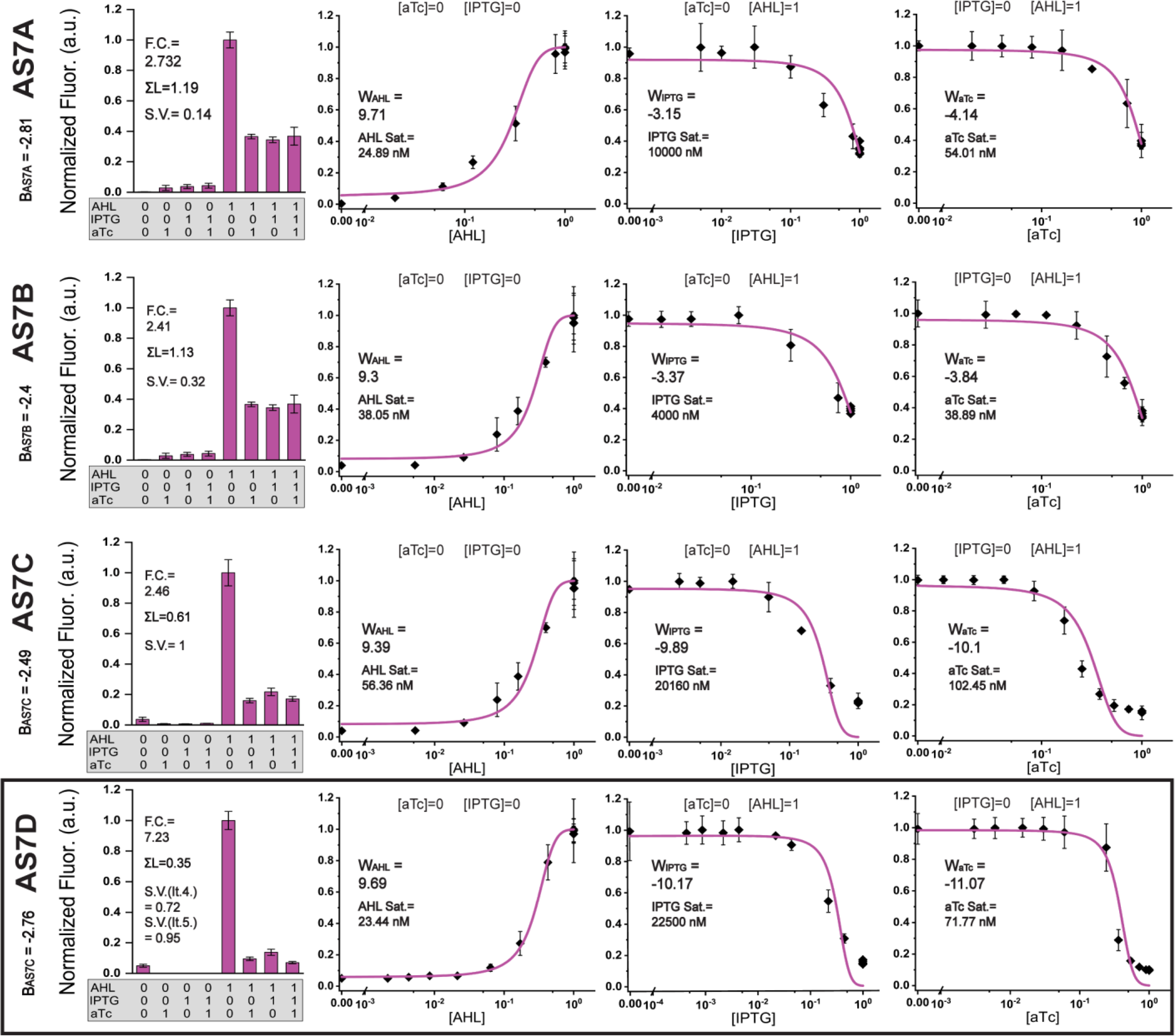
Details of characterization and Dose Responses experiments across iterations of Bactoneuron 7. (AS7A, AS7B, AS7C & AS7D) yielding corresponding parameter values (shown in figure) of Fold Change (F.C.), Σ Leakage (ΣL), Signal variation (S.V.), Bias (B_AS#_), Weights (W_Inducer_) & Inducer Saturation points (Sat.). The fluorescence readings obtained from the dose response experiments were fitted with the Log-Sigmoid function (see Methods) to derive the Weight & Bias parameters. The induction state of the two constant inducers are specified atop the fitted curves. The final selected Construct is shown boxed.

**Figure S6:**
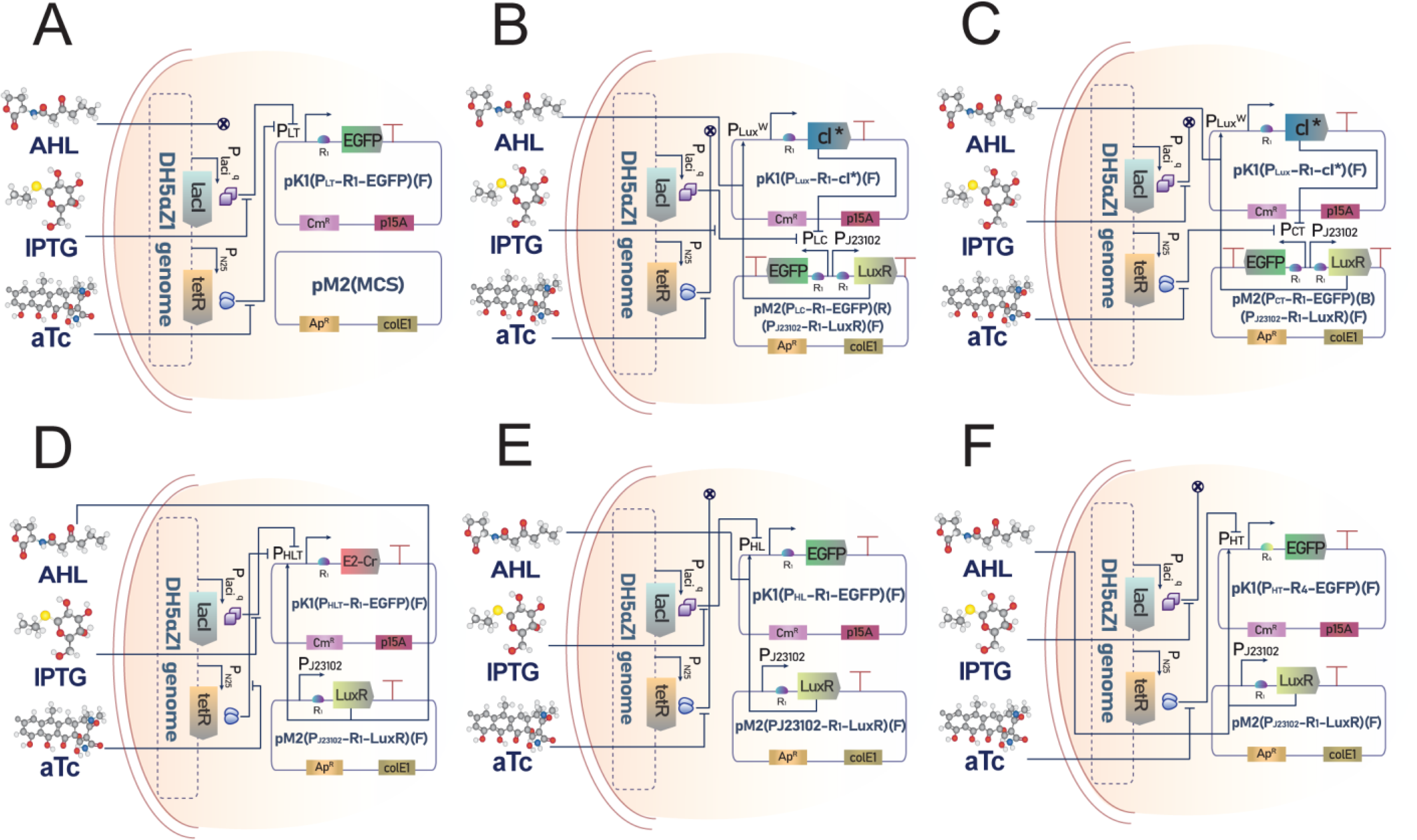
Detailed genetic circuit designs & plasmid maps of Bactoneurons that went through no iteration and were selected as final. (A) BNeu AS1, (B) BNeu AS2, (C) BNeu AS3, (D) BNeu AS4, (E) BNeu AS8, (F) BNeu AS9. Names of plasmids constructed & incorporated in the circuits are also shown within the plasmid schematic. MCS represents empty network brick cloning site. “F” & “B” denotes forward & reverse direction of cassette respectively.

**Figure S7:**
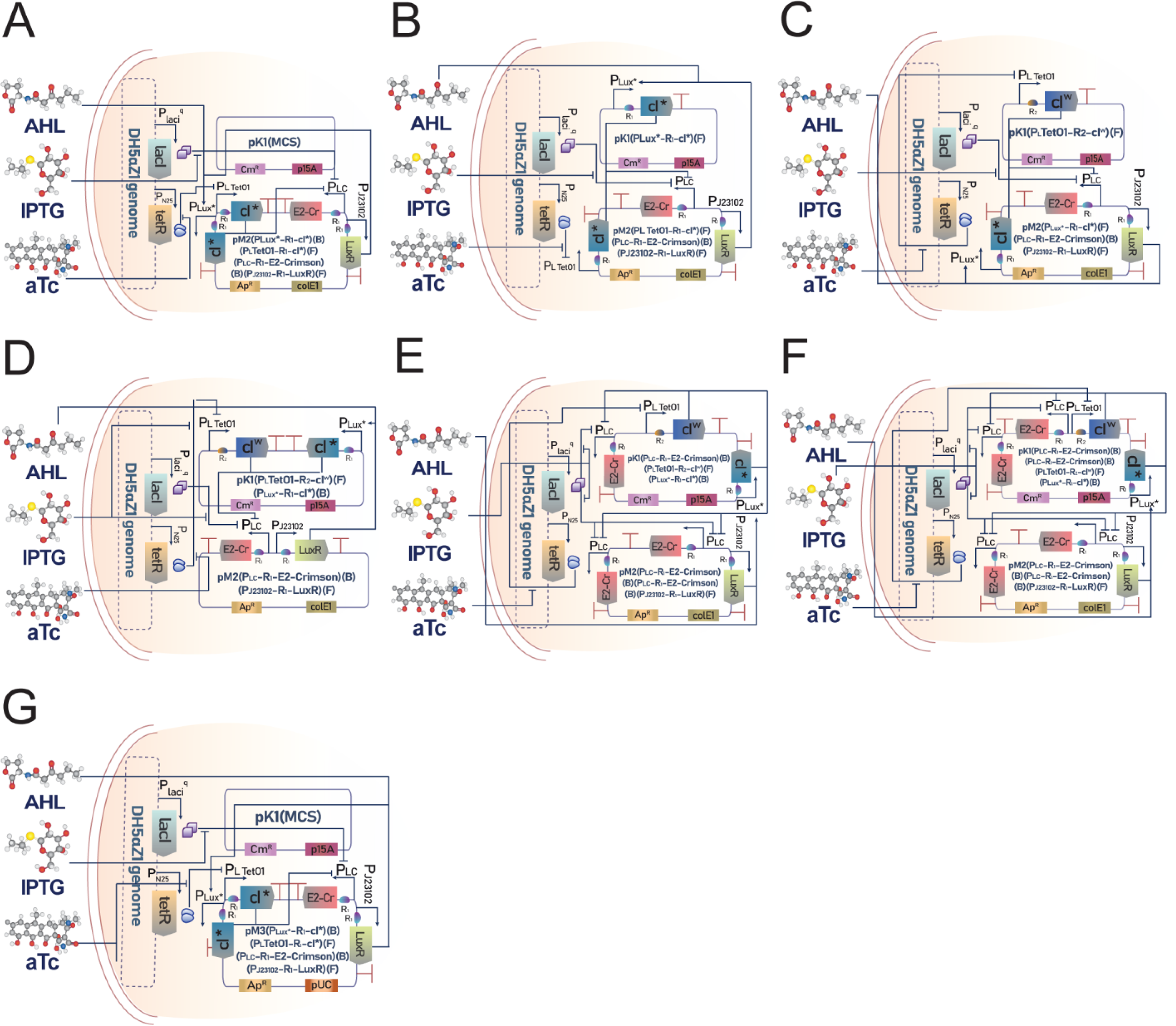
Detailed genetic circuit designs & plasmid maps of Bactoneuron 5 across iterations. (A) BNeu AS5A, (B) BNeu AS5B, (C) BNeu AS5C, (D) BNeu AS5D, (E) BNeu AS5E, (F) BNeu AS5F & (G) BNeu AS5G. Names of plasmids constructed & incorporated in the circuits are also shown within the plasmid schematic. MCS represents empty network brick cloning site. “F” & “B” denotes forward & reverse direction of cassette respectively.

**Figure S8:**
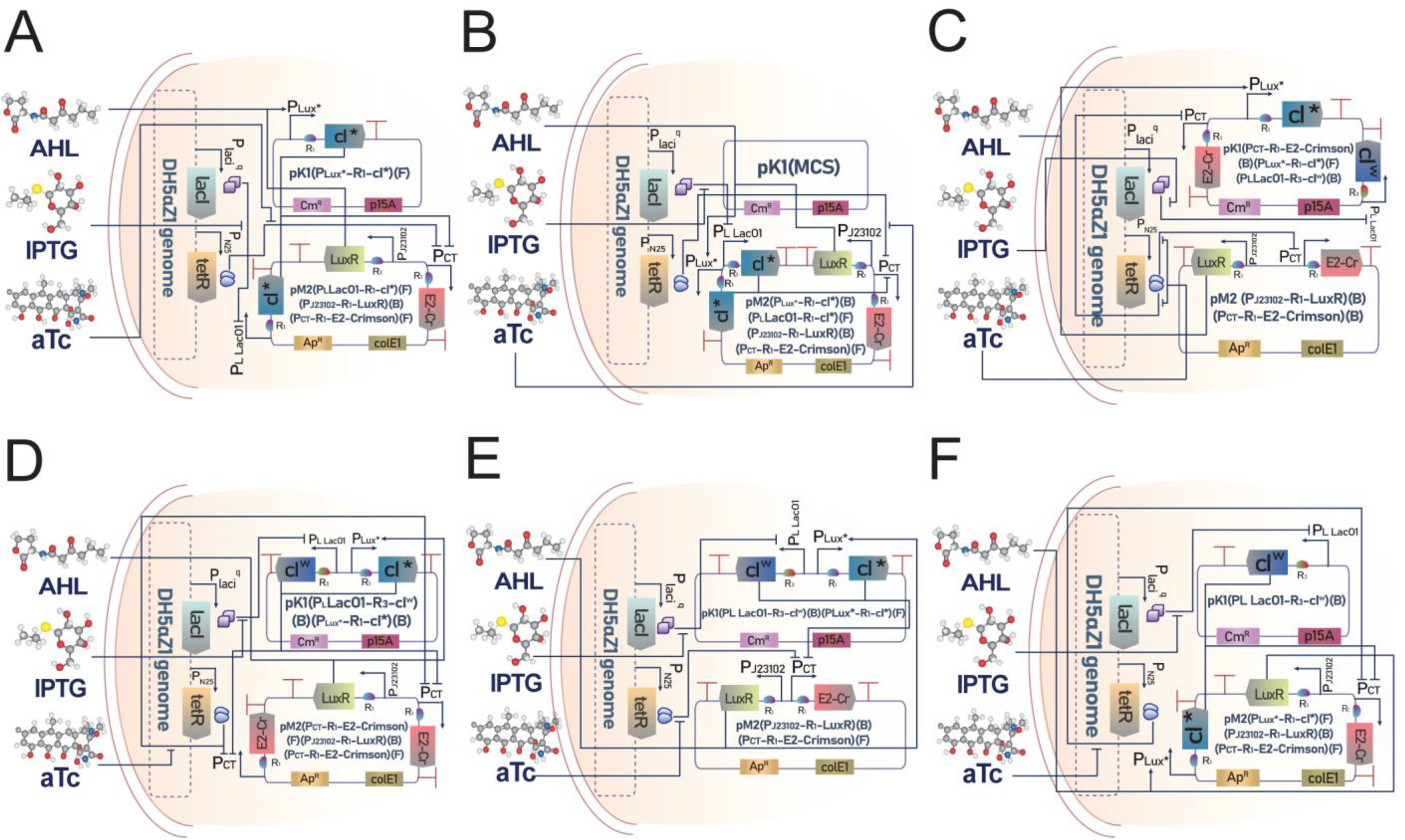
Detailed genetic circuit designs & plasmid maps of Bactoneuron 6 across iterations. (A) BNeu AS6A, (B) BNeu AS6B, (C) BNeu AS6C, (D) BNeu AS6D, (E) BNeu AS6E & (F) BNeu AS6F. Names of plasmids constructed & incorporated in the circuits are also shown within the plasmid schematic. MCS represents empty network brick cloning site. “F” & “B” denotes forward & reverse direction of cassette respectively.

**Figure S9:**
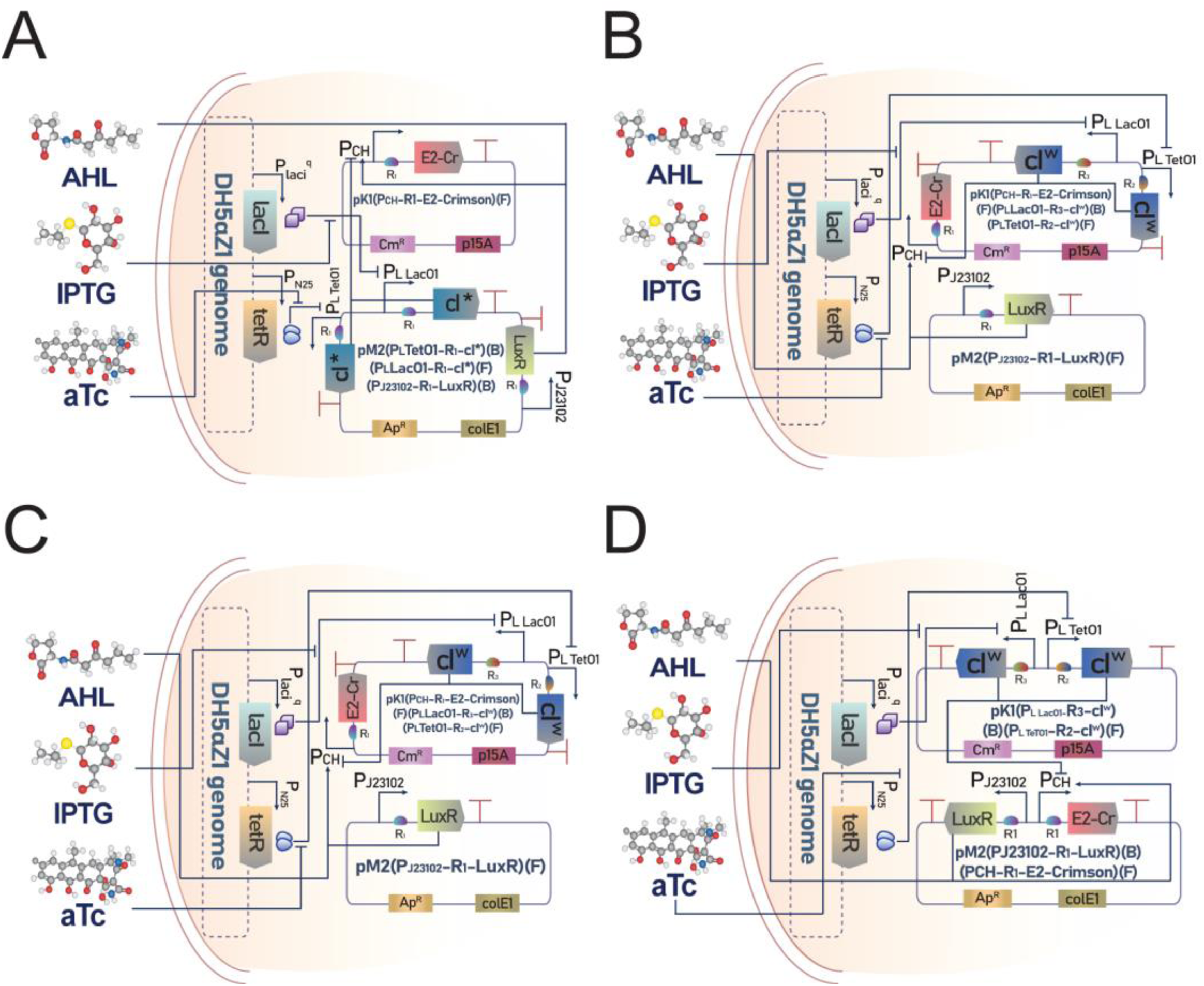
Detailed genetic circuit designs & plasmid maps of Bactoneuron 7 across iterations. (A) BNeu AS7A, (B) BNeu AS7B, (C) BNeu AS7C & (D) BNeu AS7D. Names of plasmids constructed & incorporated in the circuits are also shown within the plasmid schematic. MCS represents empty network brick cloning site. “F” & “B” denotes forward & reverse direction of cassette respectively.

**Figure S10:**
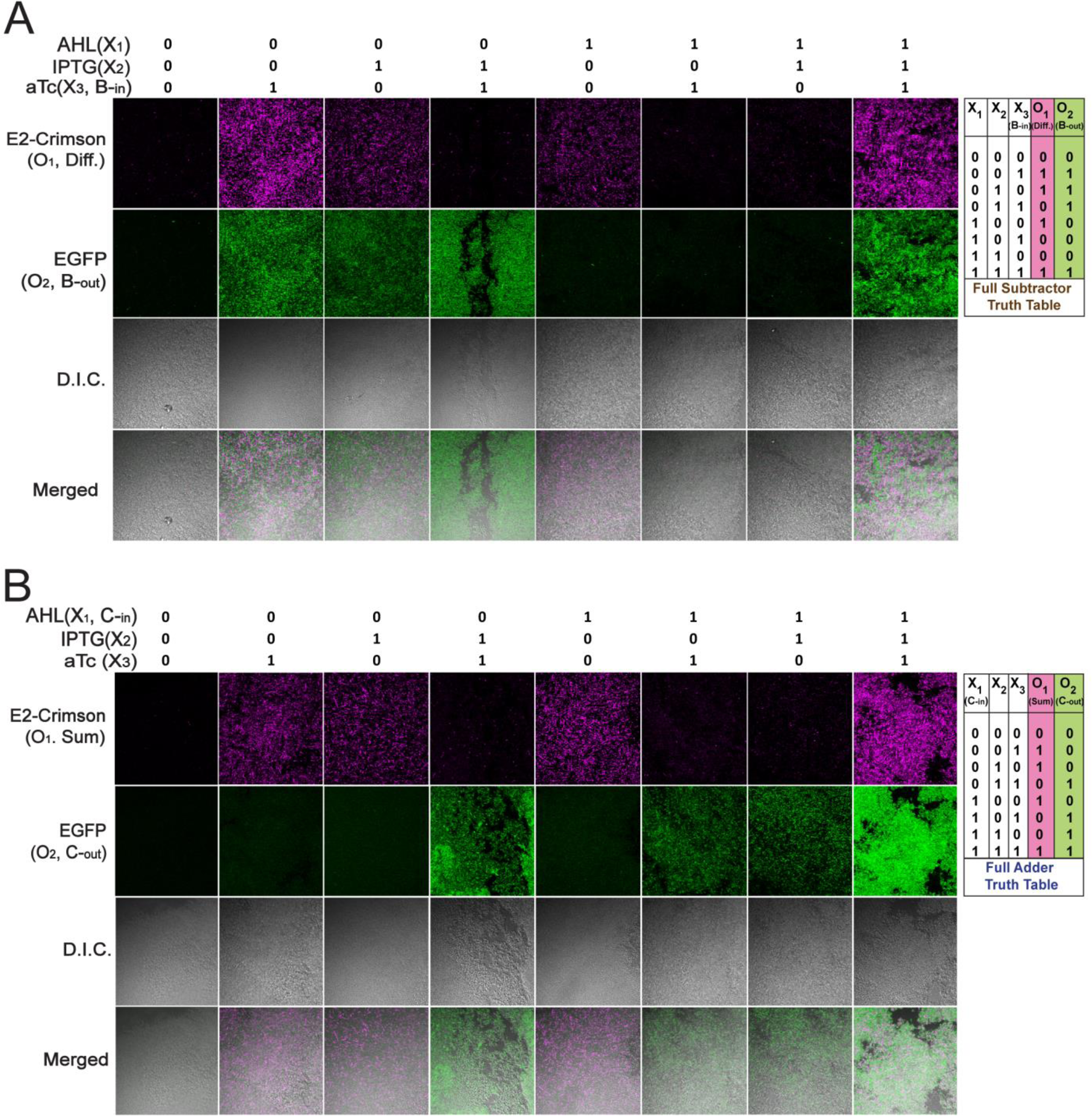
Fluorescence microscopy images with corresponding differential interference contrast (DIC) and merged channels for A) Full Subtractor, B) Full Adder. Combinations of appropriate Bactoneuron types as was required to constitute the corresponding Bactoneural layer were co-cultured for 16hours (with a sub-culture of 100 times dilution at the 10hour time point) with appropriate inducer combinations. The cultures were PBS washed and viewed as slides under relevant laser channels and emission filters. Heterogenous populations of E-coli cells forming the bactoneural layer can be seen in the fields under DIC/merged channels where they respond in accordance to the input combinations. Each sub-population of bactoneurons in the bactoneural layer either gets activated (indicated by gets fluorescence from its respective output protein) or remains inactive (showing no fluorescence) based on the input inducer combinations

**Table S1:**
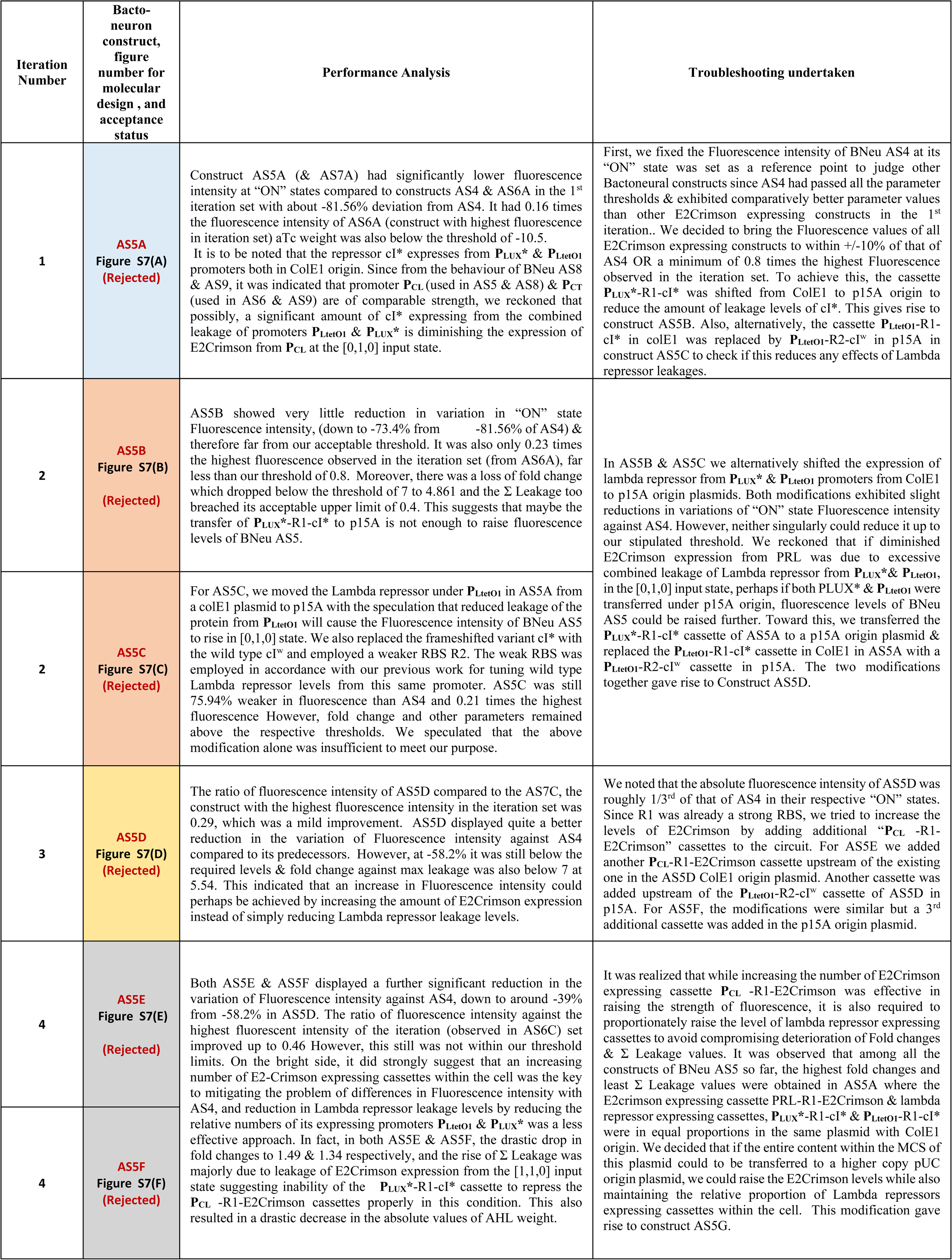
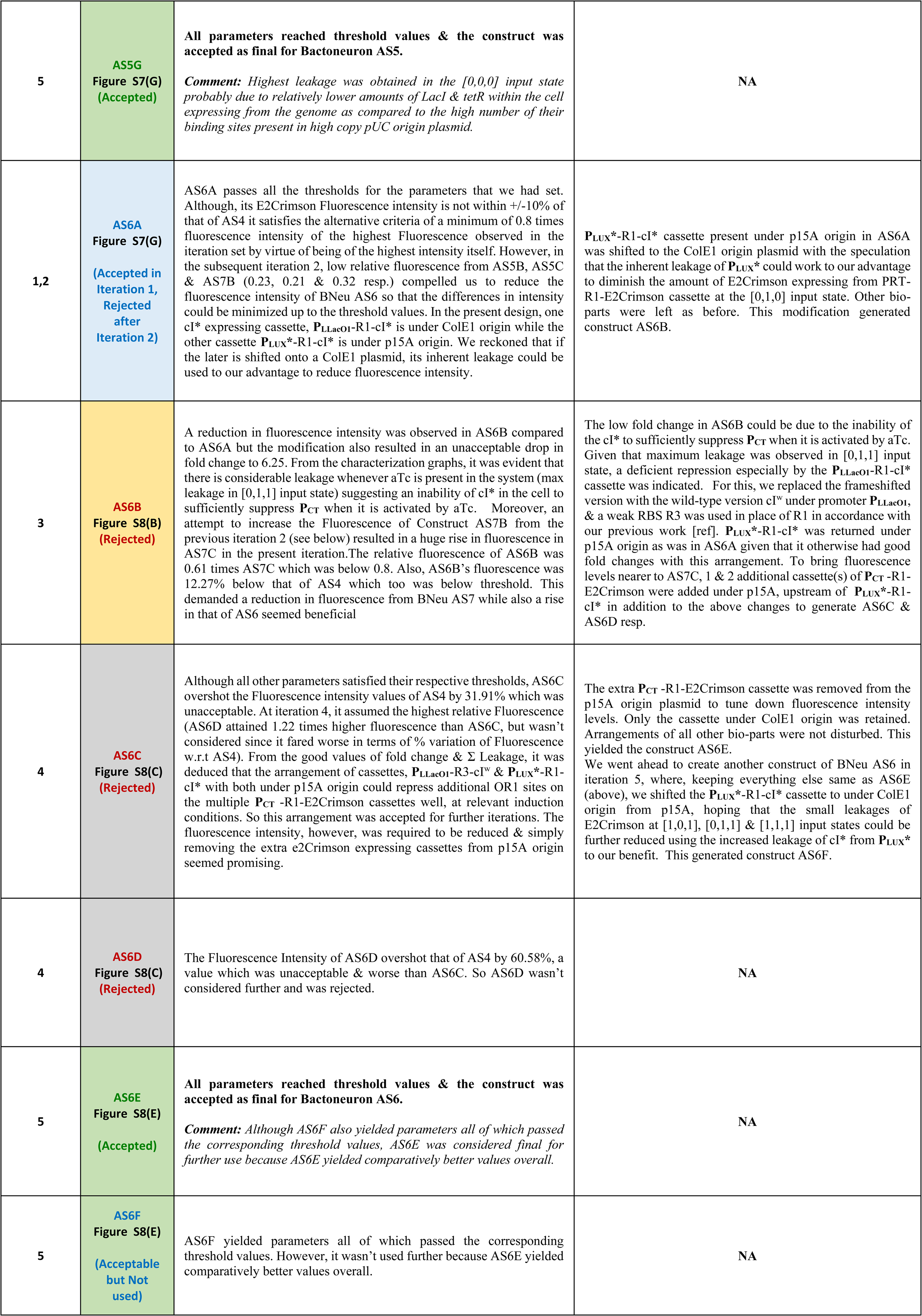
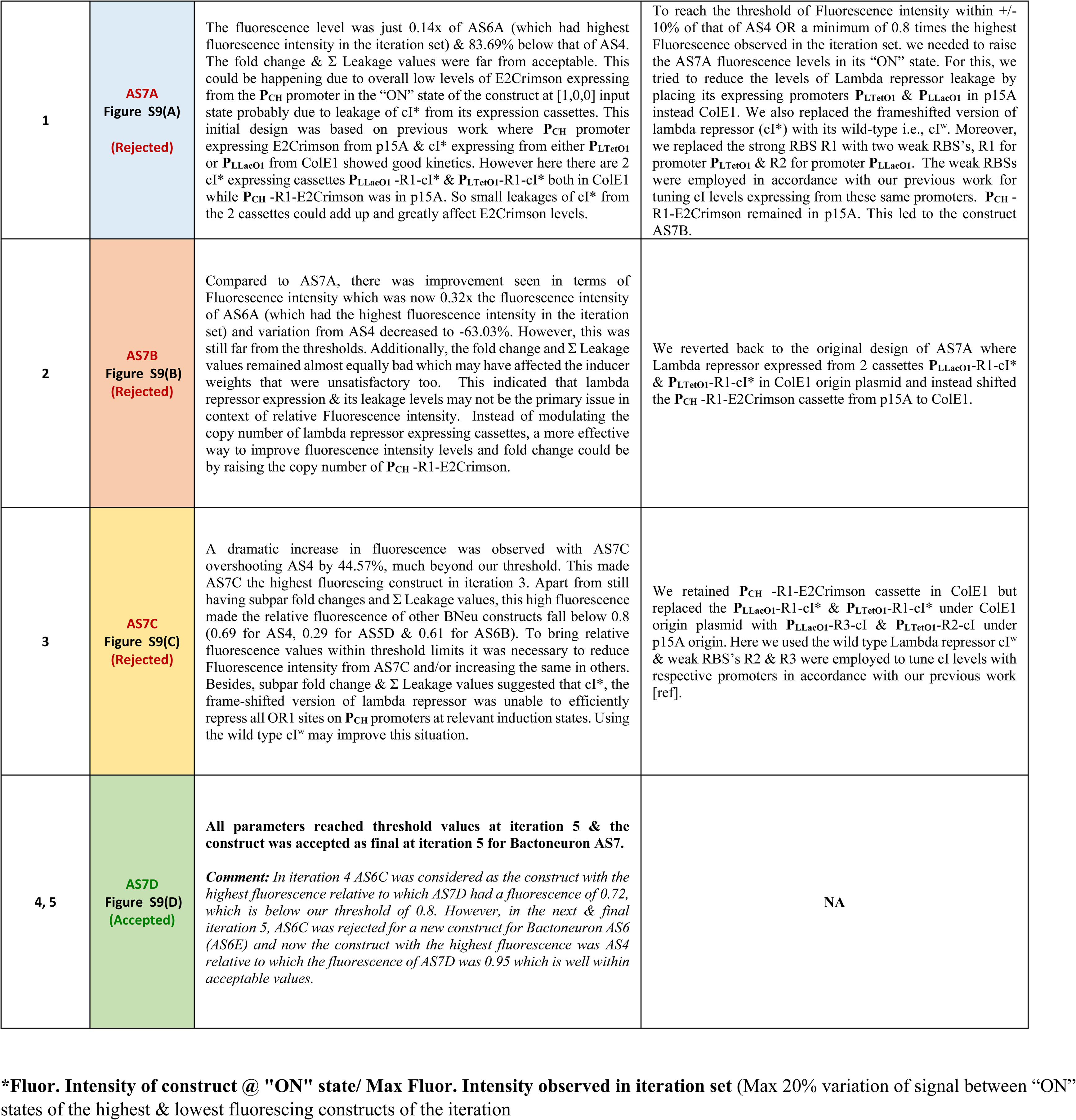
**Failure Analysis**

**Table S2:**
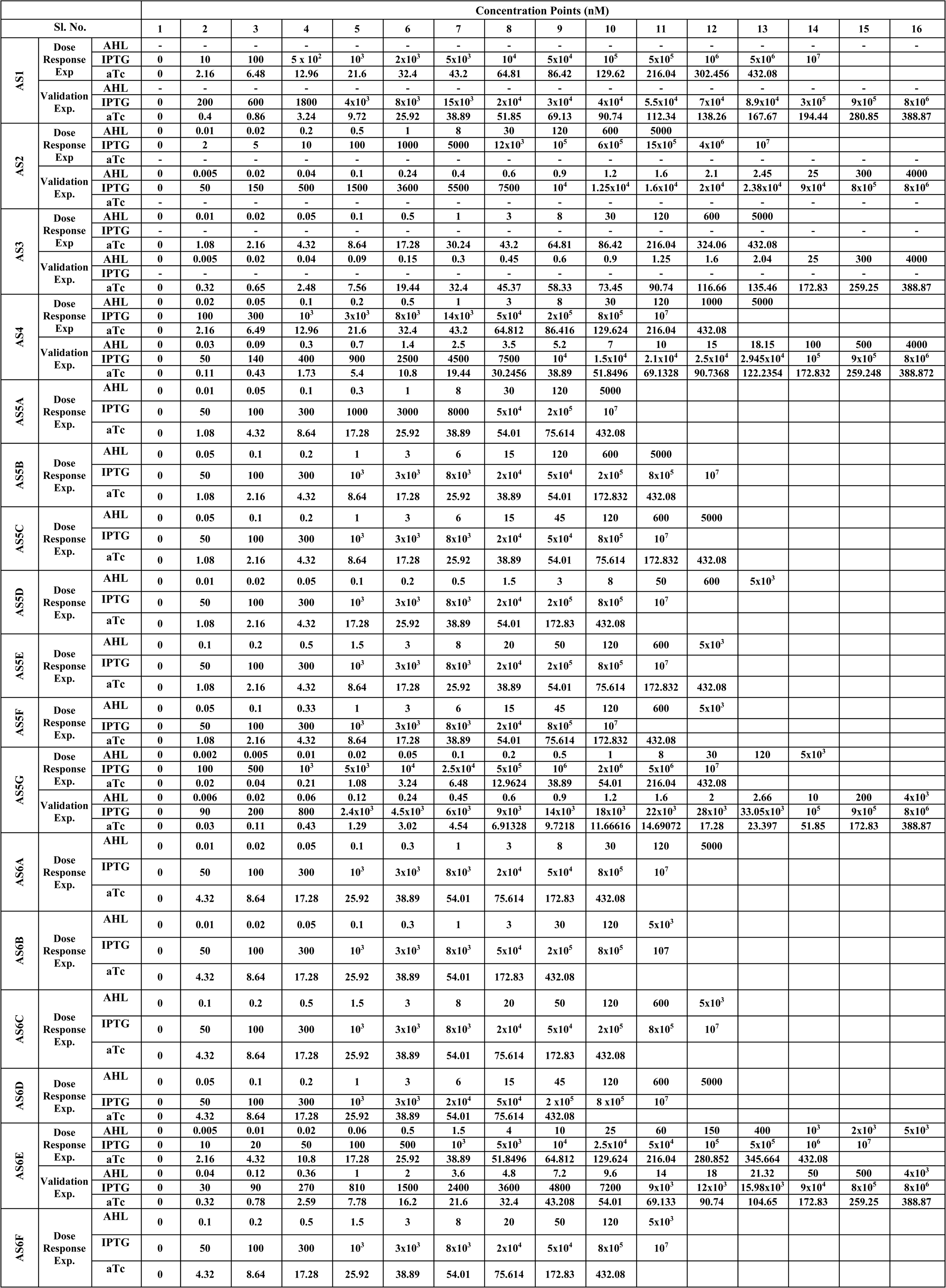
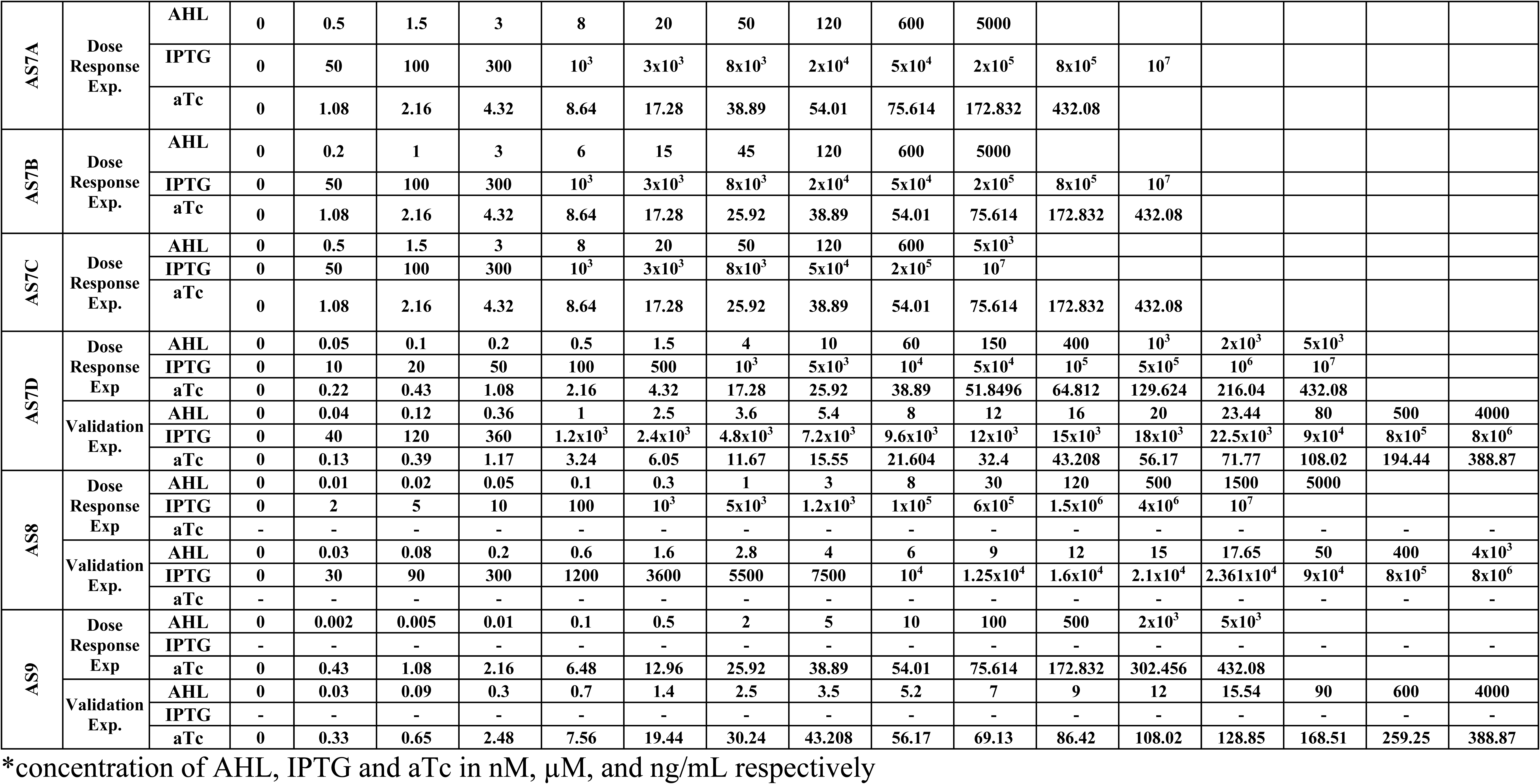
**Concentrations of inducers* used in different induction conditions for characterization, Dose response & Validation experiments of Bactoneurons**

**Table S3:**
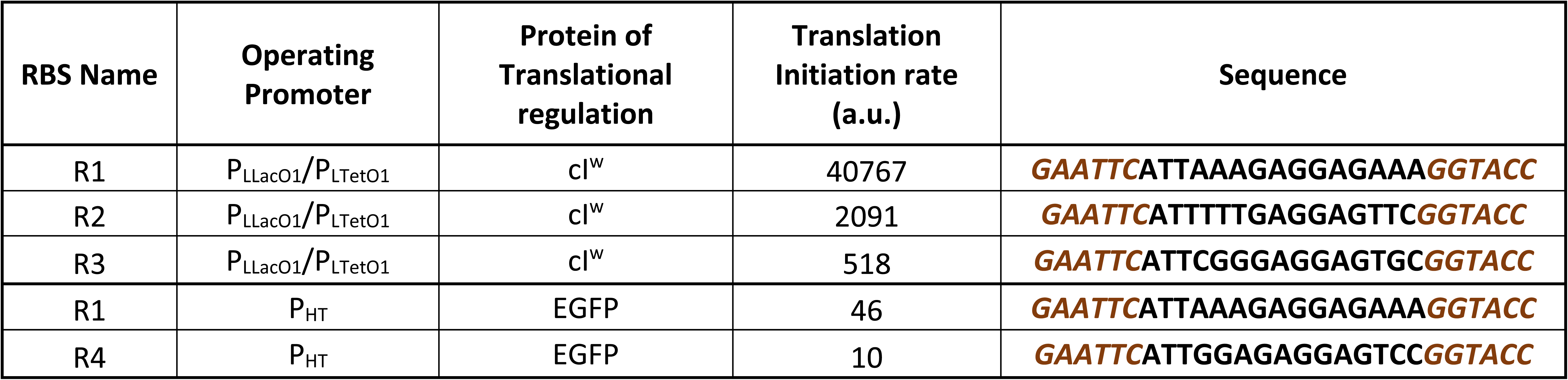
Translation initiation rate calculated from RBS calculator [32]

**Table S4:**
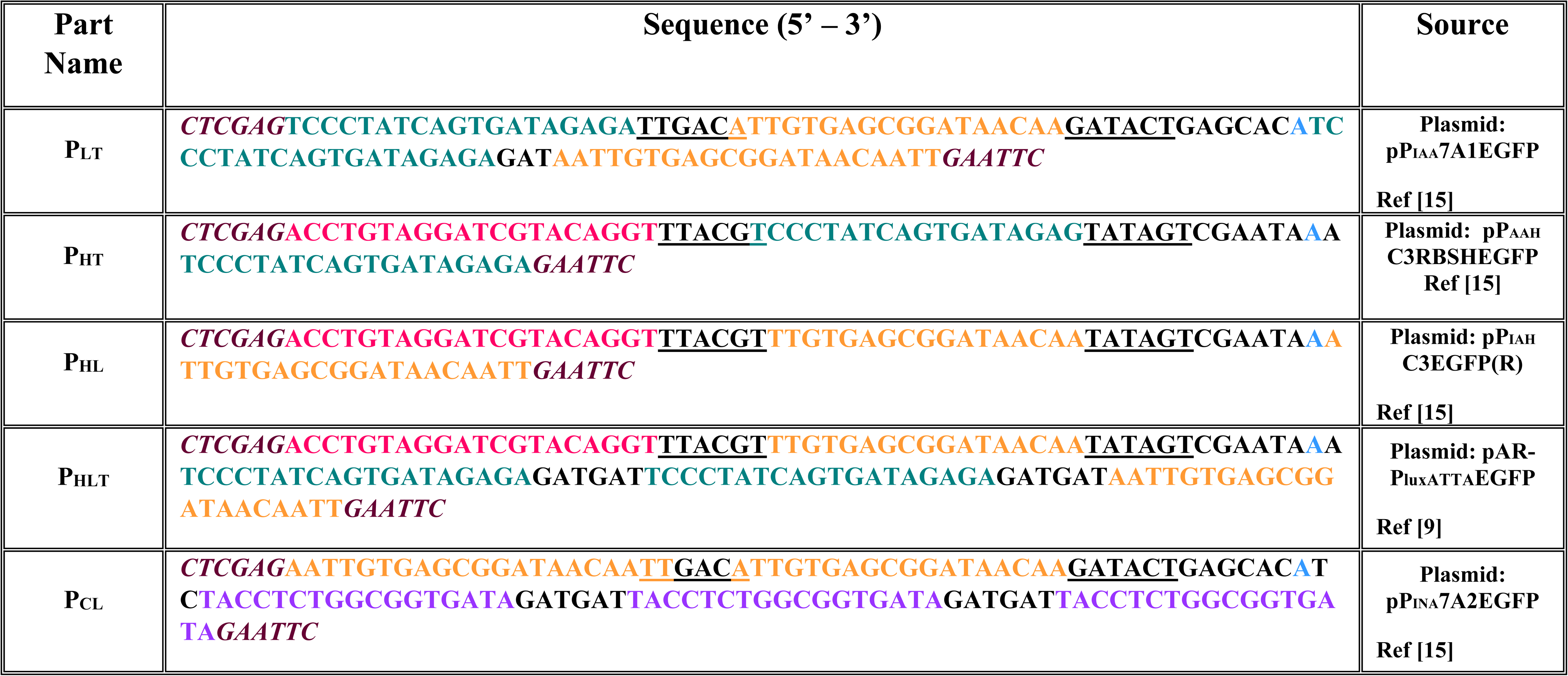
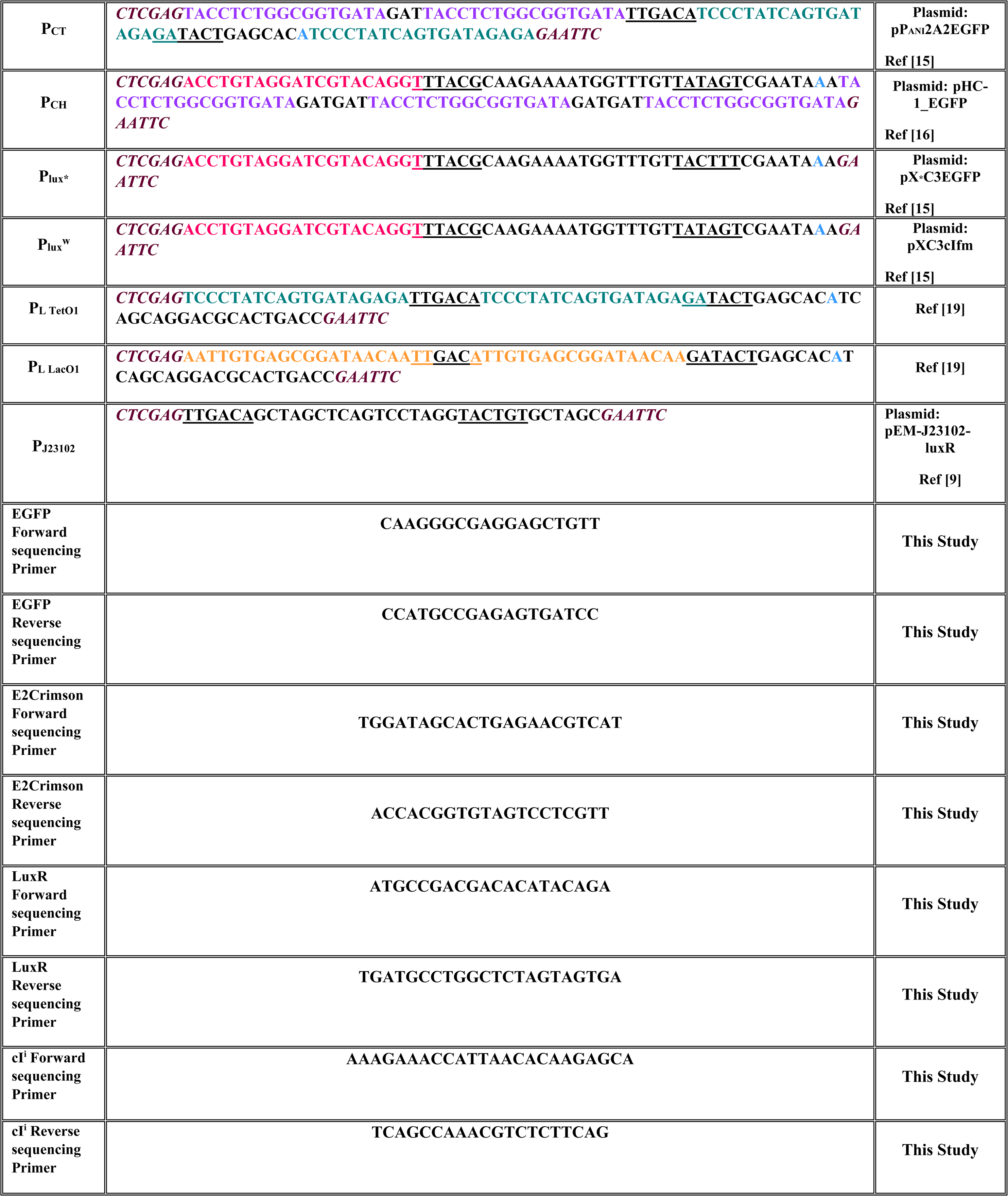
**DNA sequences of Bio-parts & primers used in this study.** lacO1, tetO2, Lux box, OR1 sites are coloured in orange, teal, pink & purple respectively. Transcription start site is shown in bold blue. -10 and -35 hexamers are underlined. Each promoter is flanked by XhoI and EcoRI restriction sites (marked in brown italics). RBS’s are flanked by EcoRI & KpnI restriction sites (marked in brown italics). Plasmids generated in this study are detailed in Figure S6, S7, S8 & S9.

**Table S5:**
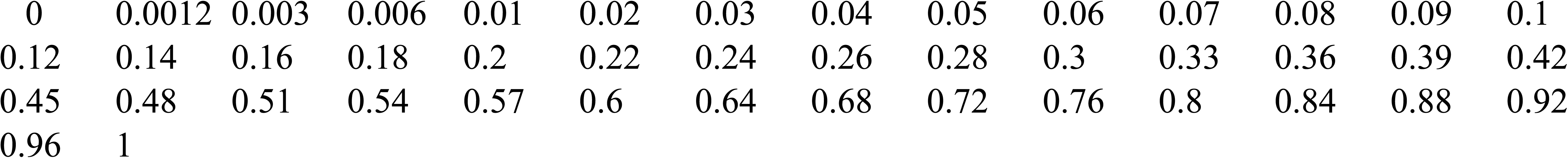
List of scaled relative concentration points to perform simulations of Bactoneurons

## References

1. Kim, M. & Julius, A.A. & Cheang U.K. Microbiorobotics: Biologically Inspired Microscale Robotic Systems: Second Edition, (Elsevier, U.K., 2017)

2. Justus KB, Hellebrekers T, Lewis DD, Wood A, Ingham C, Majidi C, LeDuc PR, Tan C.A biosensing soft robot: Autonomous parsing of chemical signals through integrated organic and inorganic interfaces. Sci Robot. 4, 88 (2019).

3. Grozinger L, Amos M, Gorochowski TE, Carbonell P, Oyarzún DA, Stoof R, Fellermann H, Zuliani P, Tas H, Goñi-Moreno A. Pathways to cellular supremacy in biocomputing. Nat Commun 10, 5250 (2019).

4. Brophy, J. A. N. & Voigt, C. A. Principles of genetic circuit design. Nature Methods 11, 508–520 (2014).

5. Chen, Y. Y., Galloway, K. E. & Smolke, C. D. Conceptual frameworks for biological design Synthetic biology: advancing biological frontiers by building synthetic systems. Genome Biology 13 240 (2012).

6. Bonnet, J., Yin, P., Ortiz, M. E., Subsoontorn, P. & Endy, D. Amplifying genetic logic gates. Science 340, 599–603 (2013).

7. Wong, A., Wang, H., Poh, C. L. & Kitney, R. I. Layering genetic circuits to build a single cell, bacterial half adder. BMC Biol 13, 40 (2015).

8. Friedland AE, Lu TK, Wang X, Shi D, Church G, Collins JJ. Synthetic gene networks that count. Science 324, 1199–1202 (2009).

9. Bonnerjee, D., Mukhopadhyay, S. & Bagh, S. Design, Fabrication, and Device Chemistry of a 3-Input-3-Output Synthetic Genetic Combinatorial Logic Circuit with a 3-Input and Gate in a Single Bacterial Cell. Bioconjug Chem 30, 3013–3020 (2019).

10. Sexton, J. T. & Tabor, J. J. Multiplexing cell-cell communication. Mol Syst Biol 16, e9618 (2020).

11. Ausländer D, Ausländer S, Pierrat X, Hellmann L, Rachid L, Fussenegger M. Programmable full-adder computations in communicating three-dimensional cell cultures. Nat Methods 15, 57–60 (2018).

12. Müller M, Ausländer S, Spinnler A, Ausländer D, Sikorski J, Folcher M, Fussenegger M. Designed cell consortia as fragrance-programmable analog-to-digital converters. Nat Chem Biol 13, 309–316 (2017).

13. Sarkar, K., Chakraborty, S., Bonnerjee, D. & Bagh, S. Distributed Computing with Engineered Bacteria and Its Application in Solving Chemically Generated 2 × 2 Maze Problems. ACS Synth Biol 10, 2456–2464 (2021).

14. Collins, J. Synthetic Biology: Bits and pieces come to life. Nature 483, S8–S10 (2012).

15. Sarkar, K., Bonnerjee, D., Srivastava, R. & Bagh, S. A single layer artificial neural network type architecture with molecular engineered bacteria for reversible and irreversible computing. Chem Sci 12, 15821–15832 (2021).

16. Srivastava, R. & Bagh, S. A Logically Reversible Double Feynman Gate with Molecular Engineered Bacteria Arranged in an Artificial Neural Network-Type Architecture. ACS Synth. Biol. 12, 1, 51–60 (2023)

17. Demuth, H. & De Jesús, B. Neural Network Design 2nd Edition, (Martin Hagan, 2014)

18. Sarkar, K., Mukhopadhyay, S., Bonnerjee, D., Srivastava, R. & Bagh, S. A frame-shifted gene, which rescued its function by non-natural start codons and its application in constructing synthetic gene circuits. J Biol Eng 13, 20 (2019).

19. Lutz, R. & Bujard, H. Independent and tight regulation of transcriptional units in Escherichia coli via the LacR/O, the TetR/O and AraC/I 1-I 2 regulatory elements. Nucleic Acids Res. 25, 1203–1210 (1997).

20. Baldwin, C. Y. & Clark, K. B. Complex Engineered Systems Ch. 9, Modularity in the Design of Complex Engineering Systems (Springer, 2006)

21. Chen, Y. Y. & Smolke, C. D. From DNA to targeted therapeutics: bringing synthetic biology to the clinic. Sci. Transl. Med. 3, 106ps42 (2011)

22. Yan, X., Liu, X., Zhao, C. & Chen, G. Q. Applications of synthetic biology in medical and pharmaceutical fields. Sig Transduct Target Ther. 8, 199 (2023).

23. Johnson, M. B., March, A. R. & Morsut, L. Engineering multicellular systems: Using synthetic biology to control tissue self-organization. Curr Opin Biomed Eng. 4 163– 173 (2017).

24. Teravest, M. A., Li, Z. & Angenent, L. T. Bacteria-based biocomputing with Cellular Computing Circuits to sense, decide, signal, and act. Energy and Environmental Science 4, 4907–4916 (2011).

25. Dvořák, P., Nikel, P. I., Damborský, J. & de Lorenzo, V. Bioremediation 3.0: Engineering pollutant-removing bacteria in the times of systemic biology. Biotechnol Adv. 35, 845–866 (2017).

26. Macia, J., Vidiella, B. & Solé, R. V. Synthetic associative learning in engineered multicellular consortia. J R Soc Interface 14, 20170158 (2017).

27. Macia, J. & Sole, R. How to make a synthetic multicellular computer. PLoS One 9, e81248 (2014).

28. Goñi-Moreno, A., Redondo-Nieto, M., Arroyo, F. & Castellanos, J. Biocircuit design through engineering bacterial logic gates. Natural Computing 10. 119–127 (2011).

29. Beal J, Goñi-Moreno A, Myers C, Hecht A, de Vicente MDC, Parco M, Schmidt M, Timmis K, Baldwin G, Friedrichs S, Freemont P, Kiga D, Ordozgoiti E, Rennig M, Rios L, Tanner K, de Lorenzo V, Porcar M. The long journey towards standards for engineering biosystems. EMBO Rep. 21, e50521 (2020).

30. Toda, S., Blauch, L. R., Tang, S. K. Y., Morsut, L. & Lim, W. A. Programming self-organizing multicellular structures with synthetic cell-cell signaling. Science (1979) 361, 156–162 (2018).

31. Aydin O, Passaro AP, Raman R, Spellicy SE, Weinberg RP, Kamm RD, Sample M, Truskey GA, Zartman J, Dar RD, Palacios S, Wang J, Tordoff J, Montserrat N, Bashir R, Saif MTA, Weiss R. Principles for the design of multicellular engineered living systems. APL Bioeng 6, 010903 (2022).

32. Mukhopadhyay, S., Sarkar, K., Srivastava, R., Pal, A. & Bagh, S. Processing two environmental chemical signals with a synthetic genetic IMPLY gate, a 2-input-2-output integrated logic circuit, and a process pipeline to optimize its systems chemistry in Escherichia coli. Biotechnol Bioeng 117, 1502–1512 (2020).

33. Salis, H. M., Mirsky, E. A. & Voigt, C. A. Automated design of synthetic ribosome binding sites to control protein expression. Nat Biotechnol 27, 946–950 (2009).

